# *In vivo* versus *in silico* assessment of potentially pathogenic missense variants in human reproductive genes

**DOI:** 10.1101/2021.10.12.464112

**Authors:** Xinbao Ding, Priti Singh, Kerry Schimenti, Tina N. Tran, Robert Fragoza, Jimmaline Hardy, Kyle Orwig, Maciej K. Kurpisz, Alexander Yatsenko, Donald F. Conrad, Haiyuan Yu, John C. Schimenti

**Affiliations:** Cornell University, College of Veterinary Medicine, Department of Biomedical Sciences, Ithaca, NY, 14853, USA; Cornell University, Department of Computational Biology, Ithaca, NY 14853, USA; Cornell University, Weill Institute for Cell and Molecular Biology, Ithaca, NY, 14853, USA; University of Pittsburgh, School of Medicine, Department of Obstetrics, Gynecology, and Reproductive Sciences, Pittsburgh, PA, 15213, USA; Division of Genetics, Oregon National Primate Research Center, Oregon Health & Science University, Beaverton, OR, 97006 USA; Fred Hutchinson Cancer Center, Seattle, WA, 98109, USA; Institute of Human Genetics, Polish Academy of Sciences, Strzeszyńska 32, 60-479, Poznan, Poland

**Author notes:** These authors contributed equally to this work (Xinbao Ding, Priti Singh).

**Keywords:** infertility, variants of uncertain significance, CRISPR/Cas9, mouse, meiosis, pathogenicity prediction algorithms, gametogenesis, reproduction

## Abstract

Infertility is a heterogeneous condition, with genetic causes estimated to be involved in approximately half of the cases. High-throughput sequencing (HTS) is becoming an increasingly important tool for genetic diagnosis of diseases including idiopathic infertility, however, most rare or minor alleles revealed by HTS are variants of uncertain significance (VUS). Interpreting the functional impacts of VUS is challenging but profoundly important for clinical management and genetic counseling. To determine the consequences of population polymorphisms in key fertility genes, we functionally evaluated 11 missense variants in the genes *ANKRD31, BRDT, DMC1, EXOI, FKBP6, MCM9, M1AP, MEI1, MSH4* and *SEPT12* by generating genome-edited mouse models. Nine variants were classified as deleterious by most functional prediction algorithms, and two disrupted a protein-protein interaction in the yeast 2 hybrid assay. Even though these genes are known to be essential for normal meiosis or spermiogenesis in mice, only one of the tested human variants (rs1460351219, encoding p.R581H in *MCM9*), which was observed in a male infertility patient, compromised fertility or gametogenesis in the mouse models. To explore the disconnect between predictions and outcomes, we compared pathogenicity calls of missense variants made by ten widely-used algorithms to: 1) those present in ClinVar, and 2) those which have been evaluated in mice. We found that all the algorithms performed poorly in terms of predicting the effects of human missense variants that have been modeled in mice. These studies emphasize caution in the genetic diagnoses of infertile patients based primarily on pathogenicity prediction algorithms, and emphasize the need for alternative and efficient *in vitro* or *vivo* functional validation models for more effective and accurate VUS delineation to either pathogenic or benign categories.

**Significance:** Although infertility is a substantial medical problem that affects up to 15% of couples, the potential genetic causes of idiopathic infertility have been difficult to decipher. This problem is complicated by the large number of genes that can cause infertility when perturbed, coupled with the large number of VUS that are present in the genomes of affected patients. Here, we present and analyze mouse modeling data of missense variants that are classified as deleterious by commonly-used pathogenicity prediction algorithms but which caused no detectible phenotype when introduced into mice by genome editing. We find that augmenting pathogenicity predictions with preliminary screens for biochemical defects substantially enhanced the proportion of prioritized variants that caused phenotypes in mice. The results emphasize that, in the absence of substantial improvements of *in silico* prediction tools or other compelling pre-existing evidence, *in vivo* analysis is crucial for confident attribution of infertility alleles.

## Introduction

A major challenge in human medical genetics is to elucidate VUS residing within disease-associated genes. In the absence of strong genetic or experimental data that validate functional consequences of such variants, namely single nucleotide variants/polymorphisms (SNVs/SNPs), functional prediction algorithms are commonly used. However, the accuracy of these predictors is not sufficiently reliable for basing clinical decisions without corroborating information (1). Compounding our relatively weak knowledge of human infertility genetics is the large number of genes that are required for normal fertility (2). Thus, even for infertile individuals who have undergone whole genome or exome sequencing (WGS and WES, respectively), an actual causal SNV or private mutation will exist within a background of VUSs in candidate genes, making it difficult to conclusively implicate any single variant as being responsible for infertility.

CRISPR/Cas9-mediated genome editing provides a means for evaluating potential human disease variants in an appropriate *in vivo* system. We previously adopted an integrated computational and experimental approach to functionally assess potentially deleterious missense variants in essential reproductive genes by modeling them in mice (3). This approach involved selection of candidate infertility missense variants based on the following criteria: a) They reside in genes that are essential for fertility in mice; b) they alter an amino acid conserved at least between mice and humans; and c) the amino acid change is predicted to be deleterious by various bioinformatic tools. However, a substantial fraction of variants tested in this manner had no clear impact on meiosis or fertility (3–6). To potentially increase the success rate in selecting actual deleterious missense variants for mouse modeling, we added two *in vitro* pre-screens to our selection pipeline (7–9). The first was to prioritize VUS that disrupt a known protein-protein interaction (PPI) since it has been shown that such human disease-causing Mendelian alleles are overrepresented (10), and the second was to identify those that destabilize the protein in cultured cells (9). These additional screens improved the yield of identifying variants that caused phenotypes (8).

Recommendations by the American College of Medical Genetics and Genomics and the Association for Molecular Pathology (ACMG/AMP) have been widely adopted by clinical laboratories around the world to guide interpretation of sequence variants (11). These guidelines provided criteria for classifying variants as pathogenic (P), likely pathogenic (LP), VUS, likely benign (LB), or benign (B). The classifications are based on distinct evidence types, each of which is assigned a level of strength. The Clinical Genome Resource (ClinGen) approves self-organized Variant Curation Expert Panels (VCEPs) in specific disease areas to make gene-centric specifications to the ACMG/AMP guidelines, and to classify variants within their scope. As a ClinGen partner, the ClinVar database archives genetic variants and assigns predicted or known impacts to diseases and other conditions (12). The collaboration of these two entities provides complementary resources to support genomic interpretation. Unfortunately, ClinGen does not have a reproductive- or infertility-related VCEP that would evaluate variants/mutations causing infertility. Consequently, compelling functional evidence from the literature becomes crucial in drawing clinical conclusions. Given the lack of appropriate or *ex vivo* systems to evaluate VUS in genes involved in certain aspects of reproduction such as spermatogenesis, and in the absence of well-characterized families segregating a suspect variant, *in vivo* evidence from knock-in/humanized animal models is currently one of the most rigorous ways to test the roles of VUS in fertility.

Here, we report experiments to test 11 missense variants in 10 genes crucial for various processes in meiosis or spermiogenesis. Nine of these are known to play essential roles in various steps of meiotic recombination or meiotic chromosome organization in mice (*Ankrd31, Brdt, Dmc1, Exo1, Fkbp6, M1ap, Mcm9, Mei1* and *Msh4*), and one (*Sept12*) is important for sperm morphogenesis and motility. We performed functional interpretation of variants in these genes by constructing and analyzing mouse models containing the orthologous amino acid substitutions. Despite *in silico* predictions (all cases) or Y2H screening (1 case) indicating that these variants are harmful, we found that mice homozygous for all but 1 of these variants - found in a male with non-obstructive azoospermia (NOA) - were fertile. Our assessment of these and other functionally interpreted alleles in mouse models, as well as infertility missense variants in the ClinVar database, reveals that informative *in vivo* assays are crucial even when computational predictions and *in vitro* assays appear compelling.

## Results

### Selection of reproduction genes and variants evaluated in this study

Primordial germ cells (PGCs), the first population of germline cells, will differentiate either toward a spermatogenic or an oogenic pathway. The process of spermatogenesis comprises multiple tightly regulated steps of differentiation (Fig. S1). It begins with several mitotic divisions of cohorts of spermatogonial stem cells that ultimately enter meiotic prophase I. During the leptotene substage of prophase I, programmed double strand breaks (DSBs) are introduced at preferred sites throughout the genome. These DSBs are recognized by a number of DNA damage response proteins that ultimately repair the DSBs via homologous recombination, and this process drives pairing and synapsis of homologous chromosomes (13). Prophase I is followed by two cell divisions without intervening DNA replication, leading to haploid, round spermatids that subsequently undergo a dramatic morphological transformation into spermatozoa during the differentiation process of spermiogenesis.

We chose to evaluate the consequences of putatively deleterious missense variants in three groups of genes (Groups 1-3) that are known to be essential for fertility in mice. These genes function in various stages of spermatogenesis as depicted in Fig. S1, and their biological roles are summarized in the following subsection. The first group of population variants (in the genes *ANKRD31, BRDT, EXO1, FKBP6, DMC1*, and *MSH4*) was selected predominantly on the bases of *in silico* prediction of pathogenicity (classified as being deleterious by the SIFT and PolyPhen2 pathogenicity prediction algorithms) and minor allele frequency (MAF) (Fig. 1). Regarding the latter, we prioritized missense variants segregating in human populations at a MAF of 0.01-1% according to the GnomAD database. Lower allele frequencies (AFs) were avoided in our original selections because cohort sample sizes available at the onset of this project were much smaller than present, raising a concern that certain “variants” were actually technical errors. Nineteen missense variants were identified under these criteria (Fig. S2A).

**Figure 1.**
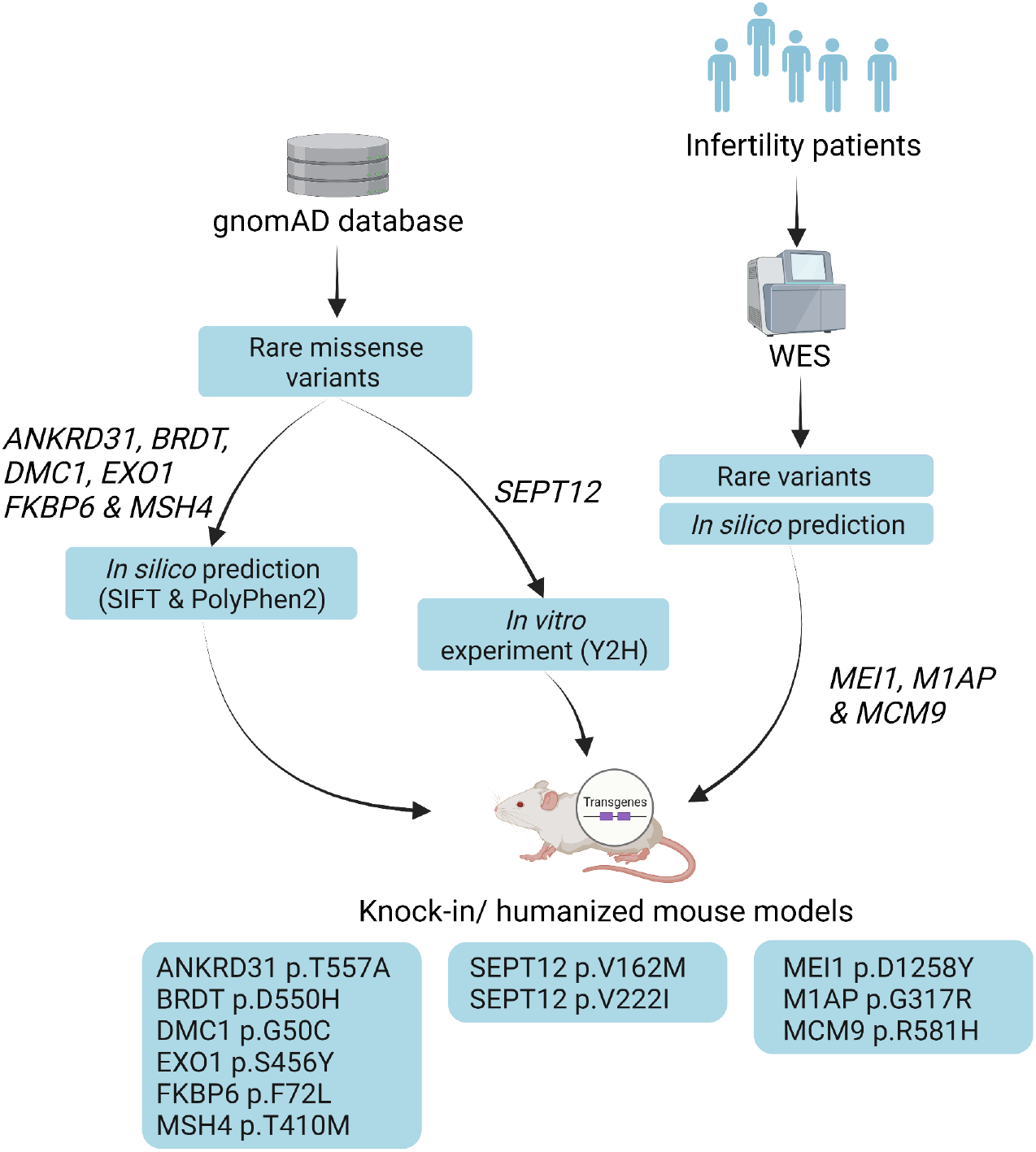
Scheme for functionally interpreting missense variants in reproduction genes. Variants were divided into 3 groups according to distinct variant prioritization pipelines.

Group 2, consisting of alleles of *SEPT12*, was selected using biochemical criteria. Mutations that disrupt PPIs are enriched as a molecular cause of human genetic diseases (10). To explore if PPI disruption might be an effective molecular screen for potential pathogenic infertility variants, yeast 2 hybrid (Y2H) assays were performed on *SEPT12* variants to determine if the altered amino acids disrupted known interactions with other Septins, SEPT1 and SEPT5 (14–16). We screened 5 variants, 4 of which disrupted a PPI (Fig. 1 and S2B). One variant, p.Gly169Glu encoded by rs138628476, was reported by our groups to cause male subfertility in a mouse model (7). The other 3 PPI-disrupting variants were added to our candidate list.

Group 3 consisted of genes (*MCM9, M1AP* and *MEI1*) harboring potentially pathogenic variants identified by whole exome sequencing of men clinically diagnosed with spermatogenic impairment (Fig.1). Like Group 1, all three alleles are polymorphisms (not private mutations) that were classified as being pathogenic by SIFT and PolyPhen2 (as well as other predictors as addressed later; Fig. 5A). The variants selected in this group have been implicated as causes of infertility in only a single case or family.

### Phenotypic analysis of mouse models

Based on our variant prioritization, Y2H screening, and resource capacity, we chose 9 missense variants from Groups 1 and 2 for functional testing by mouse modeling (Table 1). This includes 1 variant in *DMC1* that met criteria described above, except for having a lower AF than 0.01% cutoff. Additionally, the 3 variants found in infertile men were modeled in mice. Using CRISPR/Cas9-mediated genome editing in zygotes, we successfully generated 11 mouse lines modeling their corresponding human amino acid variants (Fig. S3). To identify any potential reproduction defects, the animals were bred to homozygosity and tested for several fertility parameters as described previously (3–7, 9), including testis weight, gonad histology, and sperm counts for males. For the Group 1 genes with known roles in meiosis, we also analyzed spermatocyte surface spread preparations of prophase I chromosomes via immunolabeling for key proteins. Fertility trials were performed for the relevant sex (those indicated from knockout mice) in all cases. A brief description of the genes studied, and the phenotypes of mouse models, are presented below.

**Table 1.** Variants tested in this study.

| Gene | SNP<br>(rsID) | Hs<br>AAS | Mm<br>AAS | AF (%) | SIFT | PolyPhen2 |
| --- | --- | --- | --- | --- | --- | --- |
| *ANKRD31* | rs150791065 | T557A | T544A | 0.163 | Deleterious (0.01) | Prob._damag. (1) |
| *BRDT* | rs141699970 | D550H | D546H | 0.0149 | Deleterious (0.02) | Prob._damag (0.97) |
| *DMC1* | rs189722264 | G50C | G50C | 0.0007 | Deleterious (0) | Prob._damag (0.99) |
| *EXO1* | rs4149964 | S456Y | S454Y | 0.347 | Deleterious (0) | Prob._damag (0.99) |
| *FKBP6* | rs3750075 | F72L | F72L | 0.113 | Deleterious (0) | Prob._damag (0.99) |
| *MEI1* | rs75338000 | D1258Y | D1252Y | 0.075 | Deleterious (0) | Prob._damag (0.97) |
| *M1AP* | rs140179344 | G317R | G317R | 0.007 | Deleterious (0) | Prob._damag (1) |
| *MCM9* | rs1460351219 | R581H | R581H | 0.0022 | Deleterious (0) | Prob._damag (0.99) |
| *MSH4* | rs141042002 | T410M | T432M | 0.117 | Deleterious (0.01) | Prob._damag (0.98) |
| *SEPT12* | rs144420035 | V162M | V160M | 0.049 | Deleterious (0) | Poss._damag (0.88) |
| *SEPT12* | rs142721632 | V222I | V220I | 0.189 | Tolerated (0.31) | Benign (0.29) |
| *SEPT12* | rs145805283<sup>a</sup> | T111K | T109K | 0.042 | Deleterious (0) | Prob._damag (0.96) |

AAS, amino acid substitution. AF (Allele Frequency) based on gnomAD. <sup>a</sup> No mouse model was made for this allele. Variants in *SEPT12* were evaluated by Y2H only.

DMC1 (DNA Meiotic Recombinase 1) is a meiosis-specific homolog of *E. coli* RecA and is expressed in leptotene-to-zygotene spermatocytes, stages corresponding to initiation of homologous chromosome pairing. Following generation and exonucleolytic processing of meiotic DSBs to expose 3’ overhangs bound by single stranded DNA binding protein RPA, DMC1 and a related recombinase RAD51 catalyze strand exchange, promoting homolog recognition and pairing. Male and female mouse mutants null for *Dmc1* are sterile as a consequence of failed DSB repair and interhomolog synapsis, both of which trigger prophase I arrest and checkpoint-mediated elimination of meiocytes (17, 18). We generated mice modeling the missense variant rs189722264 (NG_017203.1:g.8500G>T; p.Gly50Cys) (Fig. S3A and S4), which was predicted to be highly deleterious by both SIFT and PolyPhen2 (Table 1). In contrast to mice bearing a frameshift (presumably null) allele generated in the same editing experiment, which were azoospermic as expected, *Dmc1*^*G50C/G50C*^ female and male mice had normal fecundity (Fig. S5A). Mutant males had testis sizes and sperm counts similar to WT sibs (Fig. S5B and C), and histology revealed normal spermatogenesis, unlike the maturation arrest observed in *Dmc1*^*-/-*^ mice (Fig. S5D).

FKBP6 (FK506 binding protein 6) encodes a cis-trans peptidyl-prolyl isomerase, in which resides the amino acid altered by rs3750075 (NG_023242.2:g.6245C>A; p.Phe72Leu) (Fig. S3B and S4). This gene functions in immunoregulation and basic cellular processes involving protein folding and trafficking. Expressed in several tissues (19), FKBP6 localizes to meiotic chromosome cores and regions of homologous chromosome synapsis (20). Despite its expression in both testis and ovary, its deficiency only causes male-specific infertility (20, 21). *FKBP6* variants have been associated with NOA and idiopathic infertility (22, 23). Fertility tests of *Fkbp6*^*F72L*^ homozygotes revealed normal fecundity (Fig. S5A), testis weights, sperm counts, and testis histology (Fig. S5B-D).

EXO1 (Exonuclease 1) is involved in repair of DNA mismatches and DSBs. Loss of *Exo1* causes cancer predisposition and infertility in mice (24), the latter related to premature meiotic homolog separation before alignment at the metaphase plate (25). The SNP rs4149964 changes an amino acid residue (NG_029100.2:g.28941C>A; p.Ser465Tyr) in the MLH1 interaction domain (Fig. S3C and S4). *Exo1*^*S465Y/S465Y*^ males and females showed normal fecundity (Fig. S5A), and all male parameters were normal (Fig. S5B-D).

MSH4 (MutS protein homolog 4) is a mismatch repair protein essential for meiotic recombination. Male and female mice lacking *Msh4* are infertile due to meiotic failure (26). We modeled rs141042002 (NG_029861.1:g.56405C>T; p.Thr410Met), which is located in the DNA-binding domain (Fig. S3D and S4). The *Msh4*^*T410M/T410M*^ mutants did not reveal any reproductive phenotypes in males or females (Fig. S5A-D).

BRDT (bromodomain testis associated) belongs to the bromodomain-extra terminal (BET) family of proteins. It is expressed in pachytene spermatocytes (meiocytes in which chromosomes are fully synapsed) and round spermatids. It contains two conserved bromodomains (BD1 and 2) involved in the recognition of acetylated histones, and an extra terminal (ET) motif region involved in PPIs (27–29). Mice bearing a deletion of BD1 have oligoasthenoteratozoospermia (30). A null allele is more severe, causing complete meiotic arrest (31). As an epigenetic regulator, BRDT is required for proper meiotic chromatin organization, meiotic sex chromosome inactivation, and normal crossover formation (32).

We modeled the human *BRDT* p.Asp550His allele, encoded by SNP rs141699970, which resides in the ET domain (Figs. S3E and S4). A mouse line with a 62 nt deletion was also recovered and expanded to serve as a knockout allele (*Brdt*^*-/-*^) for comparison (Fig. S6). As previously reported (31), *Brdt*^*-/-*^ males were infertile, however, *Brdt*^*D550H/D550H*^ males sired litters of sizes comparable to those of heterozygous littermates when mated to WT females (Fig. 2A), and also had normal testis weights, sperm numbers and testis histology, in contrast to the null (Fig. 2B-D). Both *Brdt*^*-/-*^ and *Brdt*^*D550H/D550H*^ females were fertile (Fig. 2A), which was expected since *Brdt* is expressed exclusively in the testis (33). TUNEL staining revealed more apoptotic cells in the seminiferous tubules of *Brdt*^*D550H/D550H*^ mice than controls but much less than in *Brdt*^*-/-*^ mice (Fig. 2E and F). Depletion of Brdt results in weaker H3K9me3 signal on the sex chromosomes in diplonema (32). We confirmed this in *Brdt*^*-/-*^ spermatocytes but did not observe a difference between *Brdt*^*D550H/D550H*^ and control spermatocytes (Fig. 2G). However, *Brdt*^*D550H/-*^ mice exhibited slightly reduced testis weights, presence of metaphase I arrested spermatocytes, and elevated apoptotic spermatogenic cells despite having comparable sperm counts and normal H3K9me3 modification on sex body (Fig. 2B-G). Therefore, we classify *BRDT*^*D550H*^ as benign for fertility, although it may be a weak hypomorph.

**Figure 2.**
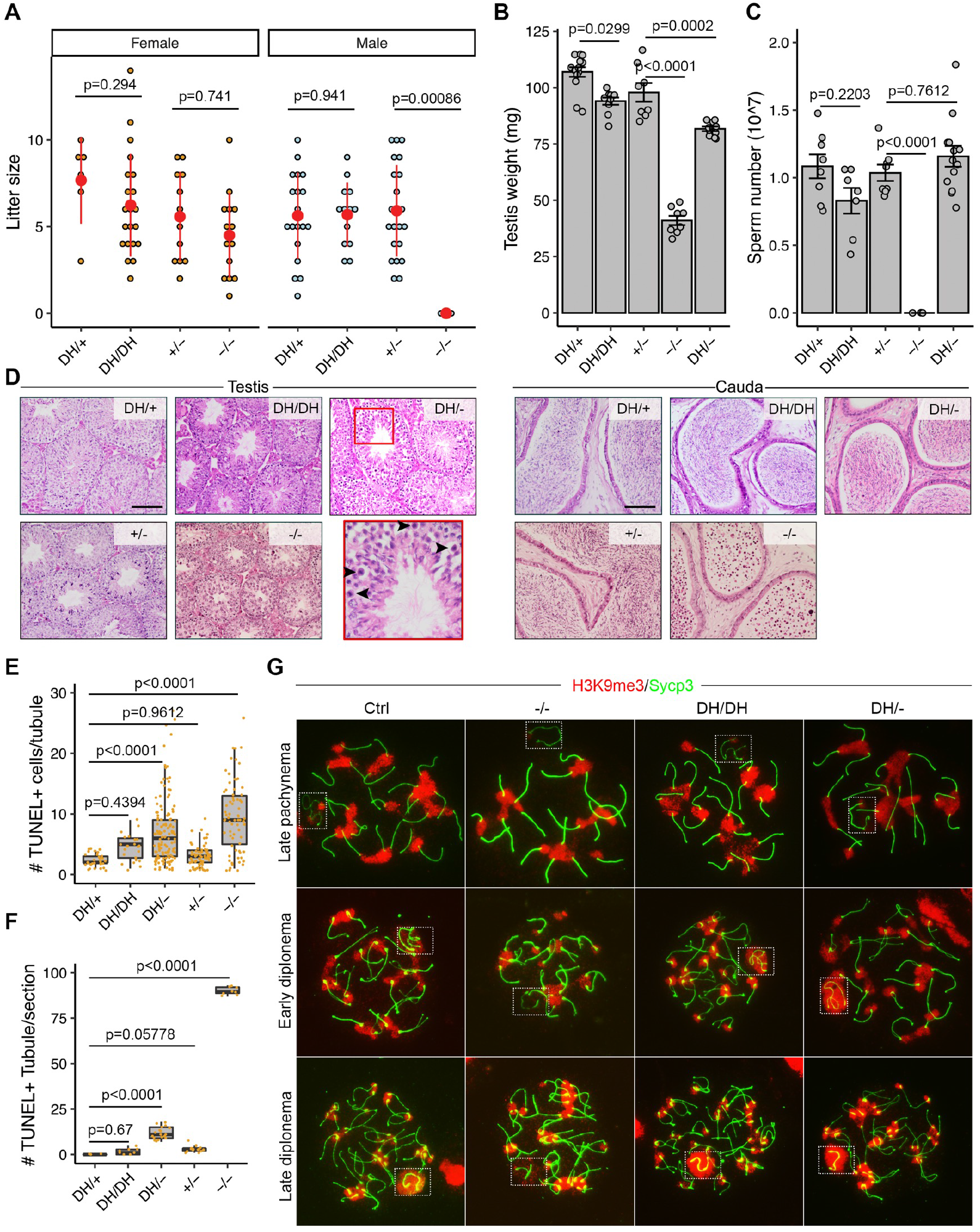
Phenotypic analysis of *Brdt*^*D550H*^ mice. A) Litter sizes from mating of *Brdt*^*D550H/D550H*^ (DH/DH), *Brdt*^*D550H/+*^ (DH/+), *Brdt*^*+/-*^ (+/-) and *Brdt*^*-/-*^ (-/-) animals to WT partners. N=2 for DH/DH male and female mice, N=3 for -/- male and female mice. B) Testis weights of 2-month-old males. C) Epididymal sperm counts (Methods). D) Histological sections of 2-month-old testes and cauda epididymides. Round germ cells were present in -/- epididymides. Scale bars = 100 μm. Black arrowheads in inset indicate meiotic metaphase I arrested cells. E) Quantification of TUNEL+ cells in each tubule. F) Number of tubules containing > 4 TUNEL+ cells. No control (DH/+) tubules section had >4. N=3 mice. G) Immunolocalization of H3K9me3 and SYCP3 on meiotic prophase I chromosomes in indicated spermatocytes stages. Dashed boxes highlight XY bodies. Data in A, B and C are represented as the mean ± SEM and were analyzed using one-way analysis of variance (ANOVA) with Tukey’s *post hoc* test.

ANKRD31 (Ankyrin repeat domain 31) contains two separated triplets of Ankyrin repeats, and three conserved domains: a predicted coiled-coil domain and two regions without functional predictions (34). Male mice lacking *Ankrd31* are sterile due to delayed recombination initiation, altered DSB distribution, and failed recombination in the pseudoautosomal region of sex chromosomes, where it is needed for DSB formation (34–36). *Ankrd31* null female mice are fertile but exhibit primary/ premature ovarian insufficiency (POI) (34, 35). We modeled human rs150791065 (NG_053151.1:g.72881A>G; p.Thr557Ala; Fig. S3F), which is located within ANK1 domain (Figs. S4). Fertility test demonstrated that *Ankrd31*^*T557A/T557A*^ mice of both sexes had normal fecundity (Fig. S7A), and homozygous males also had normal testes and sperm counts (Fig. S7B-D). Nevertheless, since *Ankrd31* is essential for recombination between sex chromosomes (34), we next examined metaphase chromosomes for this possible defect in the mutants. Chiasmata were present between each pair of homologs and between sex chromosomes in the *Ankrd31*^*T557A/T557A*^ spermatocytes (Fig. S7E). Finally, because null females are fertile despite having a reduced primordial follicle pool (34, 35), we quantified follicle numbers in 3- and 12-wk-old mutant ovaries but found no difference compared to controls (Fig. S7F and G).

SEPT12 belongs to the Septin GTP-binding protein family and is exclusively expressed in testis. It localizes around the manchette and the neck region of elongated spermatids, as well as the annulus of mature sperm. Missense mutations in *SEPT12* have been implicated as causing sperm defects and infertility in men (37). The phenotype of mice lacking *Sept12* is not clear. The first reported attempt to create mouse mutants did not obtain germline transmission, but most chimeras bearing a null allele proved to be infertile, showing a variety of testicular phenotypes (38). It is uncertain whether these phenotypes were related to haploinsufficiency for *Sept12*, or unrelated defects in the embryonic stem cells in which the targeting was performed. However, mice bearing a phosphomimetic allele caused subfertility associated with decreased sperm count and sperm motility (38).

The humanized alleles described thus far were selected through algorithmic predictions classifying them as likely pathogenic with respect to protein function. Additionally, all the alleles were selected from available population data of persons unassociated with any phenotype. We attempted to increase the likelihood that selected alleles would be consequential using two approaches. The first approach was to select variants that were pretested for a functional defect. In particular, we focused on variants that disrupt a known PPI. Accordingly, 4 alleles of *SEPT12* were selected with AF < 1% and that also disrupted at least 1 PPI. We previously performed mouse modeling of one such allele (*SEPT12*^*G169E/G169E*^) that indeed caused subfertility and poor motility (7). We failed to generate a mouse model for another (Table 1), but did generate and characterize two mouse alleles, *Sept12*^*V160M*^ and *Sept12*^*V220I*^, corresponding to human rs144420035 (NG_030315.1:g.9563G>A) and rs142721632 (NG_030315.1:g.9998G>A), respectively (Fig. S3G). Both mutations are located within a GTPase domain (Fig. S4). Males homozygous for both alleles were fertile with normal fecundity (Fig. 3A), testis weights, sperm counts, and testis histology (Fig. 3B-D). Thus, only 1 of 3 with a PPI disruption proved to be pathogenic.

**Figure 3.**
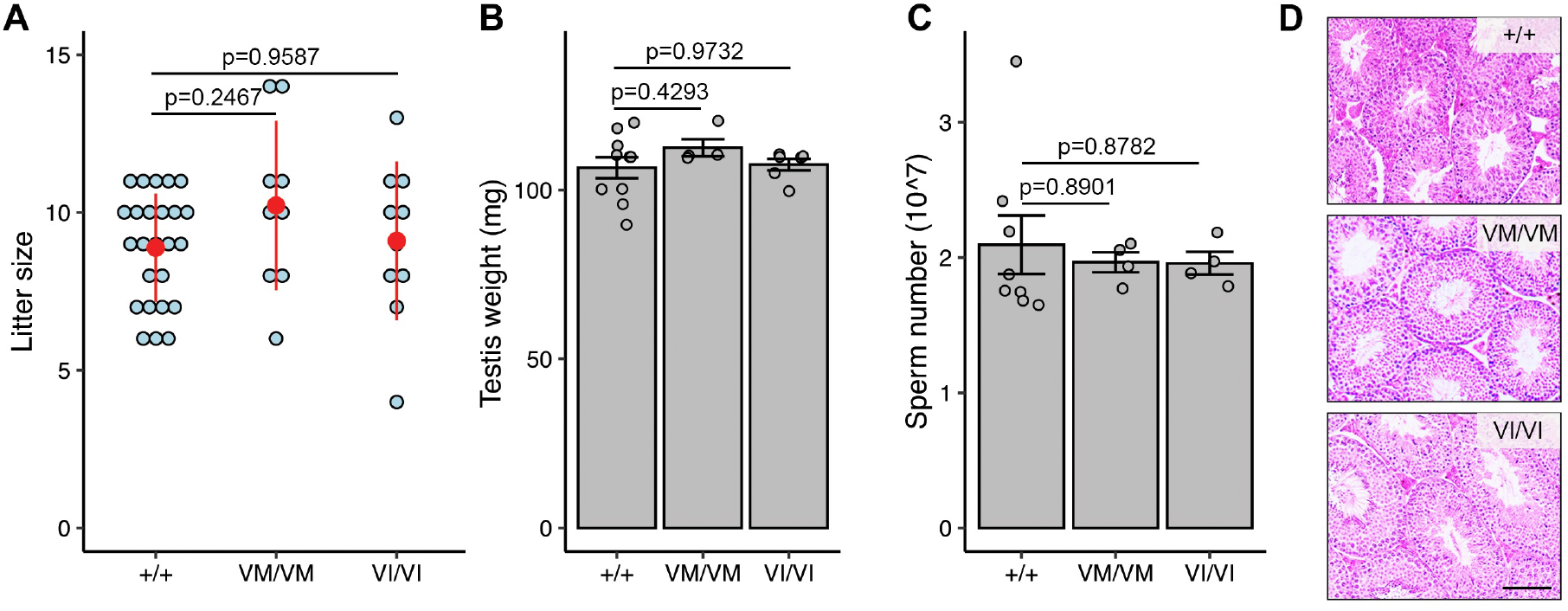
Phenotypic analysis of *Sept12* ^*V162M*^ and *Sept12*^*V222I*^ mice. A) Litter sizes from mating of WT (+/+), *Sept12*^*V162M/V162M*^ (VM/VM) and *Sept12*^*V222I/V222I*^ (VI/VI) males to WT partners. N=2 for VM/VM and VI/VI mice. B) Testis weight of 2-month-old mice. C) Sperm counts. D) Histological analyses of 2-month-old testes. Scale bars = 100 μm. Data in A, B and C are represented as the mean ± SEM and were analyzed using one-way ANOVA with Tukey’s *post hoc* test.

The other approach was to select variants (Group 3) that were identified in NOA patients, and which also were predicted to be pathogenic. As described in Methods, these three variants were identified in research studies designed to identify novel causes of NOA in human. In these studies we performed whole-genome or whole-exome sequencing of unrelated cases of idiopathic NOA, and then applied computational and statistical filters to the resulting data to generate a short list of variants that are strong candidates to be causes of NOA. The variants that pass these filters share a core set of features: they are rare in the population (<1% AF) according to GnomAD v2.1.1, they are predicted to be damaging by one or more pathogenicity prediction algorithm, and they reside in genes with gene expression that is specific to or enriched in testis.

MEI1 (meiotic double-stranded break formation protein 1) was originally identified in in a forward genetic screen in mice for infertility mutations, and was found to be required for meiotic DSB formation (39–41), a process essential for meiotic recombination and normal meiosis (42, 43). Both male and female null mouse mutants are sterile, exhibiting complete meiotic arrest from failure to synapse homologous chromosomes (39). Bi-allelic mutations of *MEI1* have also been implicated as being causal for recurrent hydatidiform moles and NOA in a family (44). Variant rs75338000 (NG_068430.1:g.162G>T) which results in the substitution of conserved amino acid Asp1258Tyr was observed in a patient with NOA due to late meiotic arrest. This variant was observed in conjunction with a 3 bp c.868_870del in-frame deletion. Both male and female *MEI1* p.Asp1258Tyr homozygotes were fertile (Fig. S3H and S4). Male sperm counts, testis weights and histology were unremarkable (Fig. S8).

M1AP (meiosis 1 associated protein) is a conserved protein of unknown biochemical function but when knocked out in mice, causes male (but not female) infertility associated primarily with meiotic metaphase I arrest (45). Bi-allelic mutations have also been associated with NOA and infertility in men (46, 47). We identified a missense variant (rs140179344; p.Gly317Arg) in a man with idiopathic NOA and histological phenotype of maturation arrest. Most *in silico* tools (7/10) classified this allele as being pathogenic, though ClinVar classified it as likely benign (Table 1; Fig. 5). We modeled this allele in mice (Fig. S3I and S4). Both male and female homozygotes were fertile. Testis weights were slightly lower in mutants, but sperm counts were not significantly reduced and testis histology appeared normal (Fig. S8). Thus, this allele does not cause NOA in mice as does the null allele or presumptive causative alleles in men.

MCM9 (minichromosome maintenance 9) is a paralog of the subunits of the heterohexameric MCM2-7 replicative helicase (48, 49). *Mcm9* hypomorphic mutant mice exhibited reduced germ cells at birth traceable to slowed proliferation of PGCs (50,51). Knockouts females (*Mcm9*^*-/-*^) were completely sterile, and males had greatly reduced (∼95%) sperm and exhibited meiotic recombination defects (52). *MCM9* variants in humans have been linked to POI in women (53–55). We modeled the variant rs1460351219 in mice encoding a predicted pathogenic variant p.Arg581His located within the conserved AAA^+^ ATPase domain of the protein (Figs.S3J; S4). This variant was found in a man with NOA who also had the predicted benign variant p.Gln658His, although the phasing was not identified. Homozygous (but not heterozygous) male and female mice were infertile. Males had reduced sperm counts, and testis histology revealed seminiferous tubule sections with variable degrees of spermatogenesis, possibly reflecting a reduction in PGCs compounded with meiotic arrest and apoptosis (Fig. 4A-C). Females display POI as almost no follicles were observed in 2-month-old ovaries (Fig. 4D). These phenotypes resemble those reported for a null allele (52).

**Figure 4.**
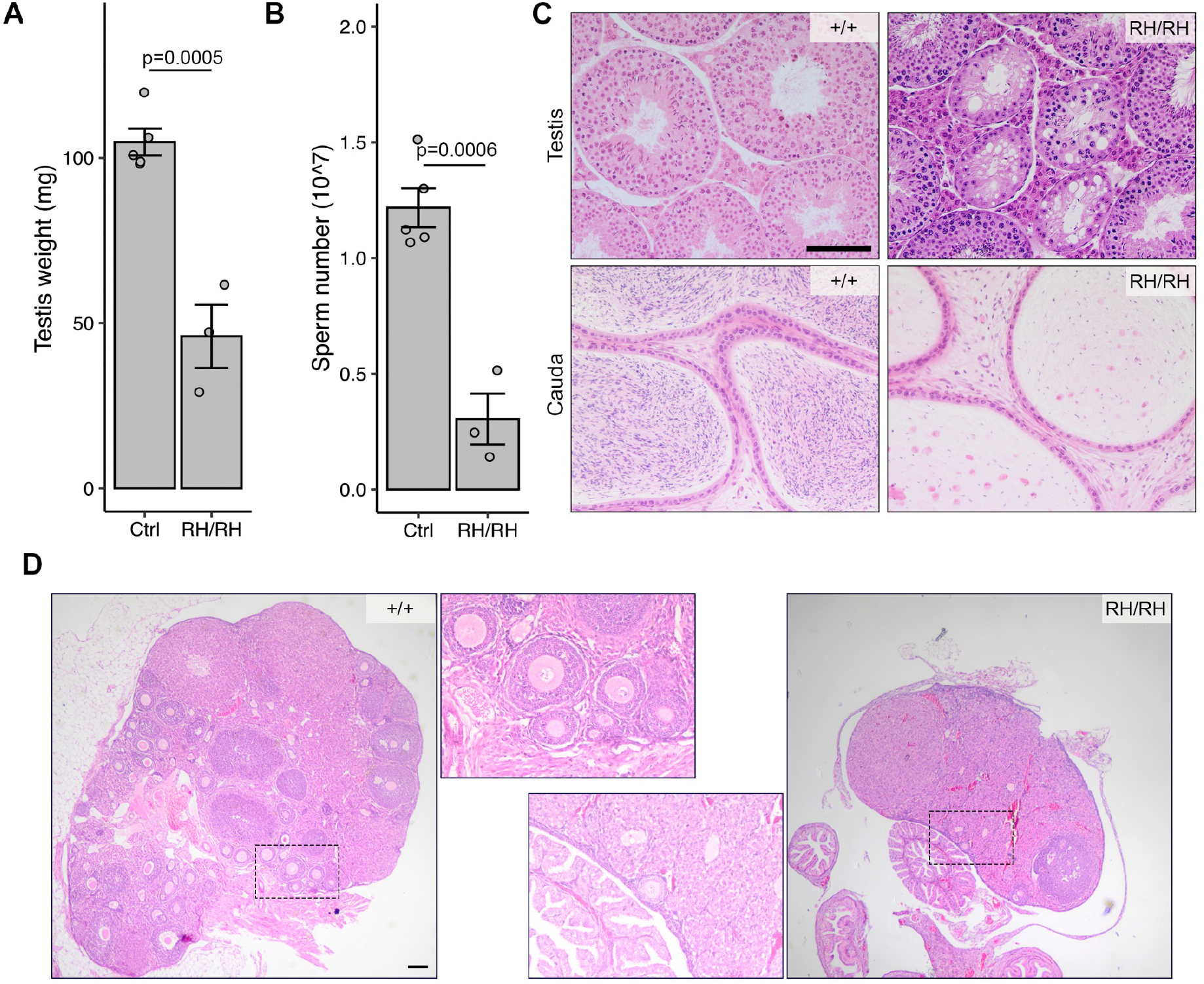
Phenotypic analysis of *Mcm9* ^*R581H*^ mice. A) Testis weights of 2-month-old mice. B) Sperm counts. C) Histological analyses of 2-month old testes. Scale bars = 100 μm. D) Representative ovary sections from 2-month old mice. The boxed regions are magnifications of follicles. Scale bars = 250 μm. Data in A and B are represented as the mean ± SEM and were analyzed using two-tailed unpaired t test.

### Performance of pathogenicity prediction algorithms in identifying disease-causing variants

To explore why such a high fraction of predicted deleterious variants failed to cause the reproductive phenotypes in mice, we evaluated the effectiveness of 10 different pathogenicity prediction algorithms in classifying 29 human alleles (including those presented here) that our group has modeled in mice, corresponding to 18 essential fertility genes (3–7, 9). We use the terms ‘deleterious’ and ‘benign’ to refer to variants that are, or are not associated with disease-related phenotypes, respectively.

Unsurprisingly, the predictors performed differently, as they use different underlying criteria. Twenty-seven of the 29 alleles were classified as deleterious by both SIFT and PolyPhen, but only 13 caused a detectable phenotype in mice. The REVEL (Rare Exome Variant Ensemble Learner) (56) and CADD (Combined Annotation Dependent Depletion; a cutoff score of >25 was used) (57) algorithms performed best by virtue of being more conservative in deleterious calls (Fig. 5A). In contrast to the low accuracy of computational predictions alone in predicting phenotypes, *in vitro* assays of variants that revealed protein defects (instability, altered function, or PPI disruptions) correctly predicted mouse phenotypes in 10/14 cases. Of the 15 alleles classified as deleterious by both PolyPhen and SIFT and for which no *in vitro* data was available, only 3 caused a mouse phenotype (Figs. 5A and S9).

**Figure 5.**
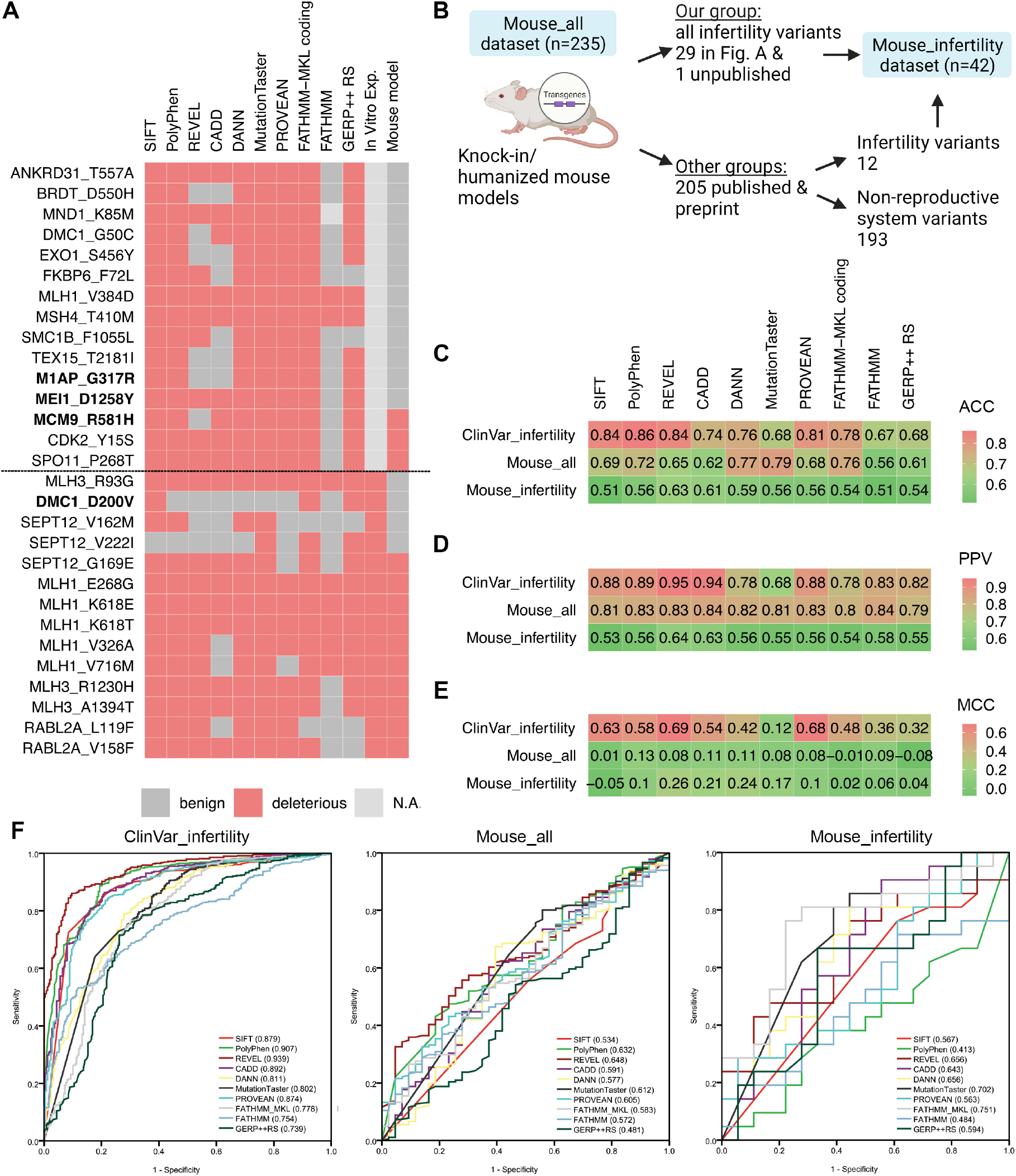
Comparison of predicted and actual deleterious variants analyzed with various pathogenicity predictors. A) *In silico* prediction outcomes of 10 commonly used algorithms for 29 functionally interpreted variants in mouse models. The *in vitro* experiments of DMC1_M200V variants were performed by Hikiba et.al., 2008. The variants below the dotted line all were tested by at least 1 *in vitro* pre-screen experiment. The bond font represents the variants were identified from infertility patients. N.A., not available. B) Overview of how the Mouse_all and Mouse_infertility datasets were derived. For missense variants modeled in mice, only those with SNP rsIDs were considered. C) Prediction Accuracy (ACC), D) Positive Predictive Value (PPV) and E) Matthews correlation coefficient (MCC) of predictors using ClinVar_infertility (n=851, only pathogenic and benign variants were used), Mouse_all (n=235) and Mouse_infertility (n=42) datasets. F) ROC curves (Receiver Operating Characteristic curve) of 10 predictors in three datasets and the AUC (Area Under the Curve) values were labeled in brackets.

The disparity between the predicted and observed effects of these missense variants led us to further explore the better-performing algorithms. We built a dataset (“Mouse_all”) comprising 235 human missense variants that have been modeled in mice, and which reside in genes involved in reproduction or other processes (Materials and Methods; Fig. 5B; Table S2). This group was subdivided into two datasets (Mouse_all_D and Mouse_all_B, where D=deleterious and B=benign) based on whether the mouse models did or did not show a pathogenic phenotype, respectively (Fig. S10A). For a “control” dataset, we retrieved infertility-related missense variants from the ClinVar database that had been assigned into five categories: ClinVar_B (benign), ClinVar_LB (likely benign), ClinVar_VUS, ClinVar_LP (likely pathogenic), and ClinVar_P (pathogenic) (Fig. S10A). For each of these datasets, we generated prediction scores from 10 algorithms computed by dbNSFP (database for nonsynonymous SNP functional predictions) (58). The prediction score distribution patterns of SIFT, DANN, GERP++ RS and MutationTaster were similar for all categories (Fig. S10B), indicating that these tools failed to discriminate the variants functionally. Next, we focused on the ClinVar_B and ClinVar_P datasets (n=851 combined, collectively referred to as “ClinVar_infertility”; Fig. S10A), representing the extremes of ClinVar classifications. Whereas the predictions of all the tools corresponded well with the pathogenic (ClinVar_P) dataset, only REVEL and CADD had relatively low FP (type I error) rates (8.43% and 8.98%, respectively) for the benign (ClinVar_B) variants. MutationTaster performed the worst (FP rate of 93.17%; Fig. S10C). Surprisingly, all of the prediction tools had higher FP rates (44.19% to 93.33%) for the Mouse_all_B group than those in ClinVar_B (infertility) group (Fig. S10C), consistent with a previous study showing that prediction tools have high FP values (59).

We next compared the performance of pathogenicity predictors for the Clinvar_infertility and Mouse_all datasets in terms of prediction accuracy (ACC), positive predictive value (PPV), Matthews Correlation Coefficient (MCC) and Area Under the Curve (AUC) of receiver operating characteristic (ROC). Consistent with previous comparisons using ClinVar variants (60), REVEL was superior to other predictors for the ClinVar_infertility dataset (ACC=0.84, PPV=0.95, MCC=0.69 and AUC=0.939; Fig. 5C-F). However, all the predictors performed worse for the Mouse_all vs the ClinVar_infertility dataset (Fig. 5C-F). The MCC scores for the Mouse_all dataset approached 0, indicating that the mouse phenotype and *in silico* interpretations are uncorrelated (Fig. 5E). We next focused on a “Mouse_infertility” data subset containing 42 functionally tested variants from the Mouse_all dataset (Fig. 5B). All the predictors performed similarly between Mouse_all and Mouse_infertility datasets (Fig. 5C-F).

As AF was a criterion for prioritizing missense variants in this study, we compared overall AF and Popmax Filtering of variants for predictive value in these three datasets. The latter method works under the assumption that there is a maximal credible population AF above which such an allele would not be compatible with disease-specific pathogenicity (61). Whereas Popmax and AF values clearly distinguished between benign vs pathogenic variants in the ClinVar_infertility dataset (Fig. 6A and Fig. S11), this was not the case for the Mouse_all and Mouse_infertility datasets (Fig. 6A and Fig. S11).

**Figure 6.**
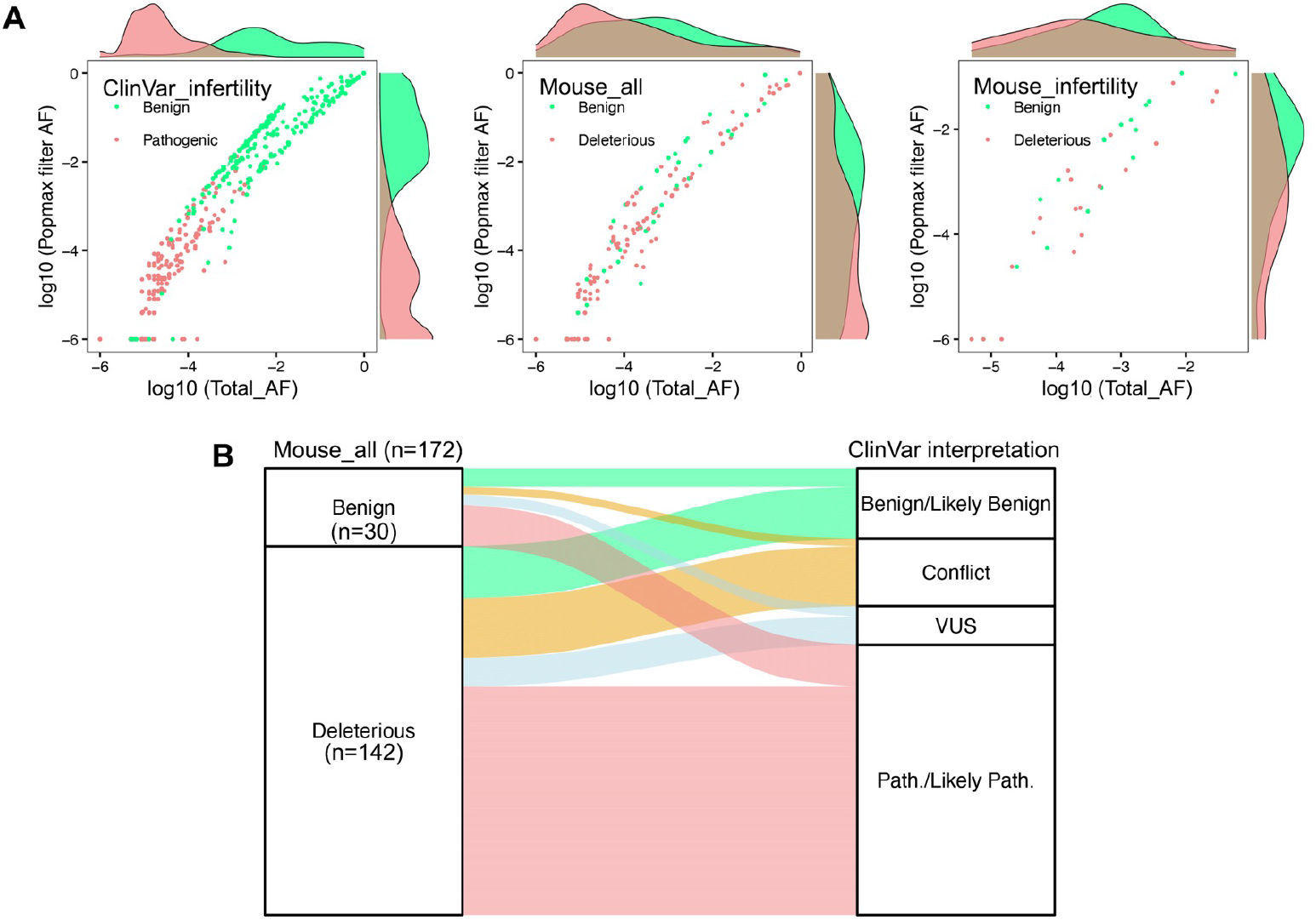
Comparison of allele frequency and variant interpretation. A) Distribution of missense variants from 3 datasets comparing total AF vs Popmax Filtering AF (95% confidence). B) Classification of functionally interpreted missense variants in ClinVar. Variants modeled in mice (n=172 interpreted in ClinVar database) were classified as either Benign or Deleterious according to the reported phenotype description. These variants were correlated with ClinVar classifications (Benign/ Likely Benign, Conflicting interpretations, VUS and Pathogenic/ Likely Pathogenic).

To study why the predictor performances differed between the ClinVar_infertility and Mouse_all datasets (*in vivo*-tested variants), we analyzed correlations between variant classifications. We retrieved ClinVar annotations of the Mouse_all dataset and found that 172 of the 235 variants (73.2%) had been interpreted in ClinVar and 55.2% (95/172) were classified concordantly (Fig. 6B). This included 16 of 45 Mouse_all benign variants being classified as “Pathogenic/Likely Pathogenic” in ClinVar, and 20 deleterious variants being classified as “Benign/Likely Benign” (Fig. 6B). With the caveat that variants may have different effects in human vs mouse, and that some of the ClinVar annotations based on patients presenting phenotypes other than infertility (for example, *MLH1/3* variants and cancer), these results raise serious questions about clinical usage of ClinVar- or *in silico*-based classifications alone and argues that *in vivo* interpreted variants, or variants with orthogonal functional evidence, are important for accurate classifications of the phenotypic consequences of genetic variants.

## Discussion

WGS and WES are quickly becoming mainstream clinical diagnostic tools for identifying genomic causes of rare diseases and cancers (62, 63). Nevertheless, wider adoption of sequencing as a diagnostic tool for other [potentially genetic] diseases is being limited by our ability to accurately classify and interpret the consequences of genetic variants. Many *in silico* prediction tools were developed to filter and prioritize variants as being potentially causative for disease, but actual phenotype correlations are difficult to predict. Complicating factors include heterogeneity of molecular mechanisms underlying diseases, lack of a deep mechanistic understanding of most genes and proteins, and complexity of physiological systems. Here, we found that *in silico* prediction tools had high FP rates when used to predict whether a human missense variant causes a phenotypic outcome in mouse models. This shortcoming underscores the need to combine computational predictions with functional evidence from appropriate cellular or animal models in order to be useful for genetic diagnosis of disease.

In clinical settings, variant classification is a complex process involving the evaluation and interpretation of multiple pieces of evidence. *In silico* predictions represent one of many evidentiary factors considered under ACMG/AMP standards. Exclusive dependence upon *in silico* tools can cause erroneous interpretation of variants. For example, SPATA16 p.Arg283Gln (64) and MEIOB p.Asn64Ile (65) were suggested as causative variants of human globozoospermia and azoospermia, respectively, but were shown not to cause reproductive phenotypes in mice, even though the latter variant caused some meiotic defects (66,67). Besides *in silico* predictors, biochemical and *in vitro* approaches are commonly used to evaluate the impact of variants on protein function. While these approaches help to prioritize or classify variants with greater confidence when used in conjunction with *in silico* predictions, the impact of such functional variants on reproductive organs or phenotype is lacking and unacceptable for use in the clinical setting as a sole diagnostic. The *DMC1* homozygous variant p.Met200Val was reported to cause POI in an African woman (68), and a subsequent study reported that this allele impaired biochemical function (69). However, we found that the orthologous allele of this highly conserved gene did not impair mouse fertility (5). In another example of non-concordance with *in vitro* data, we found that two alleles of *SEPT12* did not markedly affect mouse reproduction despite disrupting interaction with other Septins in the Y2H assay. The frequency of variants in general population is another useful criterion to assess the potential pathogenicity. Popmax Filtering AF can inform variant clinical interpretations by using a threshold - maximum credible population MAF - that is calculated based disease-specific prevalence, heterogeneity, and penetrance, and are assumed to negatively impact viability and thus frequency in the population (61). The default metric is unlikely to be as informative for infertility alleles comprising the Mouse_all dataset since viability of people is typically not impacted. To test this, we applied area under ROC curve (AUC) analysis to compare the total AF and Popmax Filtering AF in ClinVar_infertility dataset and variants interpreted by mouse models. Popmax Filtering AF did not outperform total AF in three datasets, and both AFs performed worse for variants interpreted by mouse models than classifications in the ClinVar_infertility dataset. Collectively, our data emphasize the importance of currently used animal models in the functional validation of genetic variants involved in human infertility, and highlights well-recognized challenges behind determining the physiological effects of VUS.

Our study included 3 variants that were identified by WES of infertile men. Modeling of the *MCM9*^*R581H*^ allele caused clear-cut infertility and reproductive defects in both sexes of mice, resembling knockout reproductive phenotypes. Since the mutation was not homozygous, we conjecture that the proband’s NOA was likely caused by one of the following: 1) An undetected mutation in the second allele, perhaps outside of coding regions captured by WES; 2) Epigenetic silencing of the second allele; or 3) mutations/variants in other genes that cause a synthetic phenotype in the context of the MCM9 variant. However, it is also possible that the infertility in this patient is entirely unrelated to *MCM9*^*R581H*^, especially if the second allele was functioning normally.

We concluded that the *Brdt*^*D550H*^ variant is non-pathogenic since mouse homozygotes had normal testis weight, sperm number and fertility despite the presence of increased apoptotic cells in the seminiferous tubules. Interestingly, we found that whereas mice bearing this allele in *trans* to a null (*Brdt*^*D550H/-*^) had sperm counts comparable to controls, they exhibited a subtle increase in apoptotic and meiotic metaphase I arrested spermatocytes. The mice we used were not entirely inbred, so it is possible that heterosis suppressed the phenotype. Nevertheless, there was no inconsistency amongst replicates in terms of fertility. These results suggest that functional prediction algorithms may correctly identify missense alleles that impact the protein in some subtle way that does not result in a pathogenic phenotype. However, it should be considered that the phenotype might be substantially enhanced by other background variation in a given individual.

As part of a larger effort to identify true infertility variants segregating in people, we have used pathogenicity prediction algorithms as a screen for prioritizing missense variants. However, the work summarized here reveals the shortcomings of current predictors despite being claimed to have an accuracy ranging from ∼65% to 90% (71). Given that a variety of *in silico* algorithms/pipelines are employed by clinical labs for prioritizing potential disease-causing variants (72), it is important to identify high-confidence classifiers to minimize both false-positive and false-negative prediction rates. Here, we attempted to address this issue by evaluating the performance of predictors in classifying infertility-related missense variants deposited in ClinVar. The tools varied with respect to agreement with ClinVar annotations, presumably reflecting differences in feature sets and scoring methods employed by these tools. Usually, ensemble prediction tools or metapredictors (e.g. CADD and REVEL), which generate their predictions based on the output (scores) of other tools, are purported to have higher classification accuracies than individual tools (e.g. SIFT and PolyPhen2).

To estimate the performance of a predictor, commonly used parameters include ACC, Matthews Correlation Coefficient (MCC), sensitivity, specificity, negative predictive value, PPV, and AUC. Given that all have flaws, we took an alternative approach of considering multiple success rate descriptors simultaneously. Here, we compared variant classification performance using four metrics, MCC, ACC, PPV and AUC. Consistent with a previous study (60), the consensus results from each parameter demonstrated that REVEL outperformed other classifiers in analysis of the ClinVar_infertility dataset. However, it is important to recognize that ClinVar variants are sometimes used to train algorithms either directly (such as MutationTaster) or indirectly (e.g. REVEL, which was trained using HGMD calls and shares greater than half of ClinVar variants/mutants (73)), and conversely, contributions of pathogenicity classifications to ClinVar often utilize results of various prediction algorithms (11). Such circularities in testing and training data (use of predictor calls in ClinVar classifications, and training predictors on ClinVar) may compromise accuracy of calls (74). Hence, *in vivo* tested missense variants, which are classified according to whether the functional assay (i.e., animal modeling) has a disease-causing phenotype or not, most likely constitutes a more accurate and relevant (albeit small) testing set than *in silico* predictions.

Apart from these issues, there are other potential explanations for the poor performance of predictors in classifying alleles in the Mouse_all dataset compared to those in the ClinVar_infertility dataset. One is that a significant proportion of ClinVar classifications, which are often used for algorithm training, are inaccurate. Based on data from chemical mutagenesis studies in mice, Miosge *et al*. demonstrated that for *de novo* or rare missense mutants that were algorithmically predicted to be deleterious, nearly half were in fact neutral or nearly neutral by both phenotypic and *in vitro* assays (59). Similarly, we found that all the predictors yielded high FP values (type I error). A study of human variants in the ClinVar database provided evidence for extensive misclassifications, suggested that substantial numbers of misclassifications could be corrected by considering AF, and that input from orthogonal functional and genetic studies are crucial for improving variant classification accuracy (75). A second potential explanation is that there is a substantial disconnect between biochemical and pathogenic effects of variants. That is, living systems have robustness or redundancies that largely mask minor biochemical or structural defects of proteins, although subtle molecular defects can ultimately lead to AF decline from purifying selection (59). A third possible explanation is that the power of predictors was not sufficiently evaluated by the modest sample size of the Mouse_all dataset. This possibility should become resolved as CRISPR-based mouse modeling of human variants is increasing. We anticipate that larger datasets will contribute to improving prediction tools. Finally, despite the evolutionary conservation of all the amino acids that were investigated here, it is possible that the mouse reproductive system is more robust than humans in terms of biochemical compensation for alterations, or that the consequences only manifest themselves over longer human lifespans.

Given these potential issues that can confound the accuracy of variant classification in databases, it is important to consider the levels of manual curation. ClinVar uses a three-star rating system to represent the “Review Status” of each submission. These are: “single submitter - criteria provided” (one star); “expert panel” (three stars); and “practice guidelines” (four stars). Several ClinVar derivative platforms have been developed to facilitate exploration of, and evidence for, variant interpretations. We used ClinVar Miner to identify variants of interest for evaluating the performance of pathogenicity predictors, and selected one-star rating variants because there are no infertility-related variants with three- or four-star ratings (no VCEPs exists for infertility, precluding any variants from having these ratings). The formation of an infertility VCEP for ClinGen (https://www.clinicalgenome.org) is sorely needed to critically address the accuracy of pathogenicity calls currently present in ClinVar.

In summary, we provide *in vivo* evidence for 10 of 11 human variants in essential reproduction genes to be non-pathogenic, despite being predicted to have negative impacts upon protein function by *in silico* pathogenicity prediction algorithms, and in one case, the Y2H assay. With the caveats that mice may be more tolerant to the protein alterations than humans, or that these alleles may contribute to phenotypic defects when combined with variations in other genes in individuals, we conclude that all except the MCM9 variant are likely to be clinically benign. The results underscore the utility of functional assessments of genetic variants in genetic analysis of idiopathic disease such as infertility.

## Acknowledgments

The authors would like to thank R. Munroe and C. Abratte of Cornell’s transgenic facility for generating the mice, as well as S. Wierbowski for discussions. This work is supported by grants from the National Institutes of Health (R01 HD082568 and P50 HD096723) and contract CO29155 from the NY State Stem Cell Program (NYSTEM). X.D. was supported by a postdoctoral fellowship from the Empire State Stem Cell Fund through New York State Department of Health contract C30293GG.

## Materials and Methods

### Variant prioritization and selection

The possible pathogenic functional effects of missense variants in human *DMC1, FKBP6, EXO1, MSH4, BRDT* and *ANKRD31* were analyzed using SIFT (76) and PolyPhen2 (77). The frequency of variants is as reported in Genome Aggregation Database (gnomAD v2.1.1 for *FKBP6, EXO1, MSH4, MCM9, M1AP, MEI1, BRDT* and *ANKRD31*, and v3.1.1 for *DMC1*) (gnomeAD.broadinstitute.org) as of July 2022. The domain structures of the human proteins were drawn using Domain Graph (78). The domains were identified from the literature or PhosphoSitePlus (https://www.phosphosite.org).

### Y2H screening

Y2H screening for *SEPT12* variants was performed as previously described (7). Disrupted PPIs were identified by the following criteria: a) the mutated protein reduced growth by at least 50% relative to WT as benchmarked by twofold serial dilution experiments; b) neither WT or mutant DB-ORFs were autoactivators; and c) the reduced growth phenotype was reproduced across 3 replicates. A mutation was scored as disruptive if one or more corresponding PPIs were affected.

### Selection of genetic variants identified in male infertility patients

Likely causative gene candidates were selected from WES results for adult male participants in the Genetics of Male Infertility Initiative (GEMINI) and a study conducted at the Magee-Womens Research Institute of the University of Pittsburgh (79). Patients were diagnosed with idiopathic, non-obstructive spermatogenic failure by a qualified physician on the basis of semen parameters, hormone levels, and testes histology, when available, in accordance with American Urology Association, American Society for Reproductive Medicine, and World Health Organization guidelines (80, 81). Individuals with obstructive or known causes of male infertility (e.g. CBAVD or 47, XXY karyotype) were excluded from the study.

Three variants identified in male infertility patients were nominated for mouse modeling. One mutation in *M1AP* (ENST00000536235.1: c.G949A:p.Gly317Arg) was selected from a list of variants thought to be causative for NOA identified in a cohort of 924 unrelated cases exome-sequenced by the GEMINI Consortium (82). This list of genotypes was generated by a computational pipeline intended to identify rare monogenic causes of NOA (82). This mutation was found as a heterozygote in a single sporadic case with maturation arrest testis histology and no family history of consanguinity. This case contained one other pathogenic mutation in *M1AP* (ENST00000536235.1:c.676dupT,p.W226fs), a heterozygous frameshift insertion leading to predicted loss-of-function due to nonsense mediated decay. Thus, the presumed genetic model was bi-allelic loss-of-function due to compound heterozygosity, but the phase of the variants was not determined. No other potentially causative mutations were identified in this case. Another variant, MEI1 p. Asp1258Tyr (rs75338000, NG_068430.1:g.162G>T) was observed in a patient with NOA due to late meiotic arrest. This variant was observed in conjunction with a 3 bp c.868_870del in-frame deletion. The third variant, MCM9 p.Arg581His, encoded by rs1460351219, was in a man with NOA who was also heterozygous for a predicted benign allele (rs78791427, p.Gln658His).

### Generation of mouse models by CRISPR/Cas9

All animal usage was approved by Cornell University’s Institutional Animal Care and Use Committee, under protocol 2004-0038 to J.C.S.

Mouse models were generated by the CRISPR/Cas9 genome editing technology, as described previously (3). sgRNAs and ssODNs are listed in Table S1. In addition to amino acid substitutions, several silent mutations were introduced to prevent the mutated region from being recognized and recut by the ribonucleoprotein. Briefly, the sgRNA, ssODN, and Cas9 mRNA (25 ng/μL, TriLink) were co-injected into zygotes (F1 hybrids between strains FVB/NJ and B6(Cg)-Tyr^c-2J^/J) then transferred into the oviducts of pseudopregnant females. *Dmc1, Fkbp6, Exo1, Msh4, Brdt, Mei1, M1ap, Mcm9* and *Ankrd31* founders carrying at least one copy of the desired alteration were identified and backcrossed into C57BL/6J (or FvB/NJ in the case of *Sept12* mutants) for at least three generations. For genotyping, 4 μL of crude DNA lysate was created as described previously (83) from ear punch biopsy specimens or toes of 7- to 14-day-old mice. The primers used for genotyping the mutant mice are listed in Table S1. To distinguish between WT and SNP alleles, PCR amplicons were analyzed via Sanger sequencing or restriction enzyme digestion.

### Fertility testing

Approximately two-month-old mice were housed with opposite sex WT animals for at least 3 months to determine if they were fertile. The number of pups from each mating set and sex was recorded. After at least 3 months without offspring, the analyzed individuals were considered infertile, then sacrificed for more detailed reproductive evaluations as described in the following sections.

### Sperm counts

Cauda epididymis sperm counting was performed using a standard method in the JCS laboratory (3). One cauda epididymis per male was used for each data point.

### Histology

For the preparation of paraffin blocks, tissues were fixed overnight in Bouin’s solution at room temperature, washed in 70% ethanol and then dehydrated and embedded in paraffin as previously described (3). Six-micron sections were deparaffinized and stained by hematoxylin and eosin (H&E).

### Terminal deoxynucleotidyl transferase dUTP nick end labeling (TUNEL) staining

Testes were fixed in 4% paraformaldehyde and embedded in paraffin. Sections were deparaffinized and performed TUNEL staining using the DeadEnd™ Fluorometric TUNEL System (Promega, G3250) following manufacturer’s instructions.

### Follicle counts

Ovaries were collected from mutant females and fixed in Bouin’s solution, embedded in paraffin, serially sectioned at 6 μm, and stained by H&E. Follicle quantification was performed as described previously (84). Every fifth section was examined for the presence of the following classes of oocytes/follicles: primordial, primary, secondary, preantral and antral. For quantitative analyses, they were grouped into primordial and growing follicles (from primary to antral).

### Meiotic spread preparation and immunostaining

Meiotic prophase I surface spreads were prepared as described previously (85). Briefly, testes were removed and decapsulated in hypotonic sucrose extraction buffer and left on ice for 1 h. Tubules were chopped on glass depression slides in a bubble of 0.03% sucrose and added to slides coated in 1% paraformaldehyde. Slides were slow dried and washed in PBS with Photo-Flo (Kodak).

For staining, the slides were incubated for 1 h at RT with sterile filtered antibody dilution buffer (ADB) containing 10 mL normal goat serum, 3 g BSA, 50 μL Triton X-100 and 990 mL PBS, and subsequently incubated overnight with ADB diluted rabbit polyclonal H3K9me3 (Millipore Sigma; 07-442, 1:100) and mouse monoclonal anti-Sycp3 (Abcam; ab12452, 1:200) in a humidified chamber at 4 °C. After washing in ADB, the slides were incubated at 37 °C for 30 min with a 1:1000 dilution of secondary antibody goat anti-rabbit IgG 594 (Molecular Probes, A11012) and goat anti-mouse IgG 488 (Molecular Probes, A11001), then incubated at 37 °C for 5 min with 500 ng/mL of DAPI (4′,6-diamidino-2-phenylindole) and mounted using Vectashield antifade mounting medium (Vector, H-1000).

### Diakinesis/ metaphase I chromosome spreading

This procedure was described previously (86). Briefly, seminiferous tubules were treated by hypotonic solution (1% sodium citrate) for 30 min and minced carefully for 15 min. The cell suspension was centrifuged, and the supernatant was removed. Cells were then fixed three times in methanol: glacial acetic acid (3:1) and dropped onto slides. After drying, the cells were stained with Giemsa staining solution.

### Data preprocessing

The infertility-related variants were downloaded using the ClinVar Miner web-based tool (version: 2022-2-28) (87). Variants/mutants under terms “female infertility”, “genetic infertility”, “infertility disorder” and “male infertility” with “one-star criteria provided” were selected. The variants were organized into five catalogs (pathogenic, likely pathogenic, VUS, likely benign and benign) and only missense variants with a SNP rsID were selected. Sets of deleterious and benign missense variants that have been functionally interpreted in mouse models were collected from our experiments (http://www.infertilitygenetics.org/) and from searching the literature using PubMed, Google Scholar, bioRxiv and medRxiv in June 2022 (“Mouse_all” dataset, n=235 in total, Table S2), in which the infertility variants were grouped to form the “Mouse_infertility” dataset (n=42). The analysis was restricted to SNPs with rsIDs reported in the gnomAD database. The ClinVar annotation and star rating were also collected in Table S2. Predictor calls of missense variants from the two datasets were derived using dbNSFP v4.1a (58), which is integrated into the Ensembl Variant Effect Predictor. Only the “pathogenic” and “benign” variants in ClinVar with no conflicting interpretations were grouped into the “ClinVar_infertility” dataset (n=851). The cutoff scores for predictors are listed in Table S3. Total AF and Popmax Filtering AF (95% confidence) were retrieved from gnomAD database. When transforming to logarithmic scale, 10^−6^ was added to each value because some numbers were 0.

### Evaluation of pathogenicity predictor performances

To compare the performance of the prediction tools, we applied statistical metrics derived from a confusion matrix. We identified a correctly classified variant as a true positive (TP) if, and only if, the variant corresponded to the positive class (deleterious) and as a true negative (TN) if and only if the variant corresponded to the negative class (benign). Accordingly, a false positive (FP) is a negative variant (benign) that is classified as positive (deleterious) and a false negative (FN) is a positive variant (deleterious) classified as a negative one (benign).

The Matthews Correlation Coefficient (MCC) was used to measure the correlation between the actual class of variants and the predictions made by the classifiers:

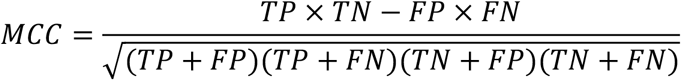

The value range of MCC is -1 to 1. A coefficient of 1 represents a perfect prediction, 0 is no better than random prediction, and −1 indicates total disagreement between prediction and observation.

Prediction precision, also known as positive predictive value (PPV), of the impact of missense variants were calculated by:

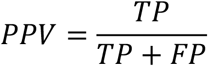

The prediction accuracy (ACC) of the impact of missense variants were evaluated using the following statistical measures:

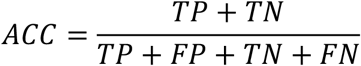

The range of values of ACC lies between 0 and 1. A perfect tool has an ACC of 1.

We used Receiver Operating Characteristic (ROC) curves to visualize the tradeoffs between sensitivity and specificity in the binary classifiers. The ROC curve is the fraction of the TP over all positives TP+FN (sensitivity) against the fraction of the FP over all negatives TN+FP (1-specificity).

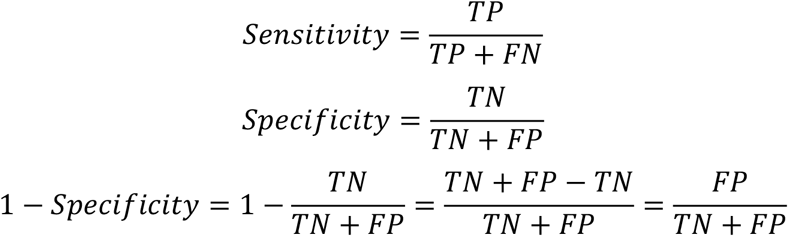

The area under the ROC curve, known as the AUC, was used to measure the performances of predictors in correctly identifying the TPs, i.e. deleterious variants, among the TPs and FPs. The AUC can take values between 0 and 1. A perfect tool has an AUC of 1 and the AUC random tool is 0.5. The prediction scores derived from dbNSFP v4.1a were used to calculate ROC in Statistical Package for the Social Sciences (SPSS) software.

### Image collections and statistics

Fluorescent images were captured by an Olympus XM10 camera. Bright-field images were captured by the Olympus SC30 camera. Cropping, color, and contrast adjustments were made with Adobe Photoshop CC 2022, using identical background adjustments for all images. All data were expressed as mean ± SEM. All statistical calculations were carried out using an unpaired Student’s *t-test* or one-way analysis of variance (ANOVA) followed by Tukey’s *post hoc* test or two-sided Fisher’s exact test with R. Graphs were generated using R. ROC curve was generated by SPSS software (version 20.0; IBM). The schematic diagrams were draw by BioRender (www.biorender.com).

## Literature Cited

1. K. Mahmood, et al., Variant effect prediction tools assessed using independent, functional assay-based datasets: implications for discovery and diagnostics. Hum Genomics 11, 10 (2017).

2. J. C. Schimenti, M. A. Handel, Unpackaging the genetics of mammalian fertility: strategies to identify the “reproductive genome”. Biol. Reprod. 99, 1119–1128 (2018).

3. P. Singh, J. C. Schimenti, The genetics of human infertility by functional interrogation of SNPs in mice. Proc Natl Acad Sci USA 112, 10431–10436 (2015).

4. T. N. Tran, J. C. Schimenti, A segregating human allele of SPO11 modeled in mice disrupts timing and amounts of meiotic recombination, causing oligospermia and a decreased ovarian reserve†. Biol. Reprod. 101, 347–359 (2019).

5. T. N. Tran, J. C. Schimenti, A putative human infertility allele of the meiotic recombinase DMC1 does not affect fertility in mice. Hum. Mol. Genet. 27, 3911–3918 (2018).

6. T. N. Tran, J. Martinez, J. C. Schimenti, A predicted deleterious allele of the essential meiosis gene MND1, present in ∼ 3% of East Asians, does not disrupt reproduction in mice. Mol. Hum. Reprod. 25, 668–673 (2019).

7. R. Fragoza, et al., Extensive disruption of protein interactions by genetic variants across the allele frequency spectrum in human populations. Nat. Commun. 10, 4141 (2019).

8. P. Singh, et al., Human MLH1/3 variants causing aneuploidy, pregnancy loss, and premature reproductive aging. Nat. Commun. 12, 5005 (2021).

9. X. Ding, et al., Variants in RABL2A causing male infertility and ciliopathy. Hum. Mol. Genet. 29, 3402–3411 (2020).

10. X. Wei, et al., A massively parallel pipeline to clone DNA variants and examine molecular phenotypes of human disease mutations. PLoS Genet. 10, e1004819 (2014).

11. S. Richards, et al., Standards and guidelines for the interpretation of sequence variants: a joint consensus recommendation of the American College of Medical Genetics and Genomics and the Association for Molecular Pathology. Genet. Med. 17, 405–424 (2015).

12. M. J. Landrum, et al., ClinVar: improvements to accessing data. Nucleic Acids Res. 48, D835–D844 (2020).

13. S. Gray, P. E. Cohen, Control of meiotic crossovers: from double-strand break formation to designation. Annu. Rev. Genet. 50, 175–210 (2016).

14. T. Rolland, et al., A proteome-scale map of the human interactome network. Cell 159, 1212–1226 (2014).

15. H. Yu, et al., Next-generation sequencing to generate interactome datasets. Nat. Methods 8, 478–480 (2011).

16. E. L. Huttlin, et al., The BioPlex Network: A Systematic Exploration of the Human Interactome. Cell 162, 425–440 (2015).

17. D. L. Pittman, et al., Meiotic prophase arrest with failure of chromosome synapsis in mice deficient for Dmc1, a germline-specific RecA homolog. Mol. Cell 1, 697–705 (1998).

18. K. Yoshida, et al., The mouse RecA-like gene Dmc1 is required for homologous chromosome synapsis during meiosis. Mol. Cell 1, 707–718 (1998).

19. X. Meng, X. Lu, C. A. Morris, M. T. Keating, A novel human gene FKBP6 is deleted in Williams syndrome. Genomics 52, 130–137 (1998).

20. M. A. Crackower, et al., Essential role of Fkbp6 in male fertility and homologous chromosome pairing in meiosis. Science 300, 1291–1295 (2003).

21. K. Shah, G. Sivapalan, N. Gibbons, H. Tempest, D. K. Griffin, The genetic basis of infertility. Reproduction 126, 13–25 (2003).

22. W. Zhang, S. Zhang, C. Xiao, Y. Yang, A. Zhoucun, Mutation screening of the FKBP6 gene and its association study with spermatogenic impairment in idiopathic infertile men. Reproduction 133, 511–516 (2007).

23. T. Miyamato, et al., Is a genetic defect in Fkbp6 a common cause of azoospermia in humans? Cell. Mol. Biol. Lett. 11, 557–569 (2006).

24. K. Wei, et al., Inactivation of Exonuclease 1 in mice results in DNA mismatch repair defects, increased cancer susceptibility, and male and female sterility. Genes Dev. 17, 603–614 (2003).

25. S. Schaetzlein, et al., Mammalian Exo1 encodes both structural and catalytic functions that play distinct roles in essential biological processes. Proc Natl Acad Sci USA 110, E2470–9 (2013).

26. B. Kneitz, et al., MutS homolog 4 localization to meiotic chromosomes is required for chromosome pairing during meiosis in male and female mice. Genes Dev. 14, 1085–1097 (2000).

27. C. Pivot-Pajot, et al., Acetylation-dependent chromatin reorganization by BRDT, a testis-specific bromodomain-containing protein. Mol. Cell. Biol. 23, 5354–5365 (2003).

28. B. D. Berkovits, D. J. Wolgemuth, The role of the double bromodomain-containing BET genes during mammalian spermatogenesis. Curr. Top. Dev. Biol. 102, 293–326 (2013).

29. J. Shi, C. R. Vakoc, The mechanisms behind the therapeutic activity of BET bromodomain inhibition. Mol. Cell 54, 728–736 (2014).

30. E. Shang, H. D. Nickerson, D. Wen, X. Wang, D. J. Wolgemuth, The first bromodomain of Brdt, a testis-specific member of the BET sub-family of double-bromodomain-containing proteins, is essential for male germ cell differentiation. Development 134, 3507–3515 (2007).

31. J. Gaucher, et al., Bromodomain-dependent stage-specific male genome programming by Brdt. EMBO J. 31, 3809–3820 (2012).

32. M. Manterola, et al., BRDT is an essential epigenetic regulator for proper chromatin organization, silencing of sex chromosomes and crossover formation in male meiosis. PLoS Genet. 14, e1007209 (2018).

33. M. H. Jones, M. Numata, M. Shimane, Identification and characterization of BRDT: A testis-specific gene related to the bromodomain genes RING3 and Drosophila fsh. Genomics 45, 529–534 (1997).

34. F. Papanikos, et al., Mouse ANKRD31 Regulates Spatiotemporal Patterning of Meiotic Recombination Initiation and Ensures Recombination between X and Y Sex Chromosomes. Mol. Cell 74, 1069-1085.e11 (2019).

35. M. Boekhout, et al., REC114 partner ANKRD31 controls number, timing, and location of meiotic DNA breaks. Mol. Cell 74, 1053-1068.e8 (2019).

36. L. Acquaviva, et al., Ensuring meiotic DNA break formation in the mouse pseudoautosomal region. Nature 582, 426–431 (2020).

37. Y.-C. Kuo, et al., SEPT12 mutations cause male infertility with defective sperm annulus. Hum. Mutat. 33, 710–719 (2012).

38. Y.-H. Lin, et al., The expression level of septin12 is critical for spermiogenesis. Am. J. Pathol. 174, 1857–1868 (2009).

39. B. J. Libby, L. G. Reinholdt, J. C. Schimenti, Positional cloning and characterization of Mei1, a vertebrate-specific gene required for normal meiotic chromosome synapsis in mice. Proc Natl Acad Sci USA 100, 15706–15711 (2003).

40. L. G. Reinholdt, J. C. Schimenti, Mei1 is epistatic to Dmc1 during mouse meiosis. Chromosoma 114, 127–134 (2005).

41. B. J. Libby, et al., The mouse meiotic mutation mei1 disrupts chromosome synapsis with sexually dimorphic consequences for meiotic progression. Dev. Biol. 242, 174–187 (2002).

42. P. J. Romanienko, R. D. Camerini-Otero, The mouse Spo11 gene is required for meiotic chromosome synapsis. Mol. Cell 6, 975–987 (2000).

43. S. Keeney, C. N. Giroux, N. Kleckner, Meiosis-specific DNA double-strand breaks are catalyzed by Spo11, a member of a widely conserved protein family. Cell 88, 375–384 (1997).

44. N. M. P. Nguyen, et al., Causative mutations and mechanism of androgenetic hydatidiform moles. Am. J. Hum. Genet. 103, 740–751 (2018).

45. N. A. Arango, et al., Meiosis I arrest abnormalities lead to severe oligozoospermia in meiosis 1 arresting protein (M1ap)-deficient mice. Biol. Reprod. 88, 76 (2013).

46. M. J. Wyrwoll, et al., Bi-allelic Mutations in M1AP Are a Frequent Cause of Meiotic Arrest and Severely Impaired Spermatogenesis Leading to Male Infertility. Am. J. Hum. Genet. 107, 342–351 (2020).

47. C. Tu, et al., An M1AP homozygous splice-site mutation associated with severe oligozoospermia in a consanguineous family. Clin. Genet. 97, 741–746 (2020).

48. B. K. Tye, MCM proteins in DNA replication. Annu. Rev. Biochem. 68, 649–686 (1999).

49. K. Yoshida, Identification of a novel cell-cycle-induced MCM family protein MCM9. Biochem. Biophys. Res. Commun. 331, 669–674 (2005).

50. Y. Luo, J. C. Schimenti, MCM9 deficiency delays primordial germ cell proliferation independent of the ATM pathway. Genesis 53, 678–684 (2015).

51. S. A. Hartford, et al., Minichromosome maintenance helicase paralog MCM9 is dispensible for DNA replication but functions in germ-line stem cells and tumor suppression. Proc Natl Acad Sci USA 108, 17702–17707 (2011).

52. M. Lutzmann, et al., MCM8- and MCM9-deficient mice reveal gametogenesis defects and genome instability due to impaired homologous recombination. Mol. Cell 47, 523–534 (2012).

53. M. A. Wood-Trageser, et al., MCM9 mutations are associated with ovarian failure, short stature, and chromosomal instability. Am. J. Hum. Genet. 95, 754–762 (2014).

54. S. Desai, et al., MCM8 and MCM9 nucleotide variants in women with primary ovarian insufficiency. J. Clin. Endocrinol. Metab. 102, 576–582 (2017).

55. T. Guo, et al., Novel pathogenic mutations in minichromosome maintenance complex component 9 (MCM9) responsible for premature ovarian insufficiency. Fertil. Steril. 113, 845–852 (2020).

56. N. M. Ioannidis, et al., REVEL: an ensemble method for predicting the pathogenicity of rare missense variants. Am. J. Hum. Genet. 99, 877–885 (2016).

57. P. Rentzsch, M. Schubach, J. Shendure, M. Kircher, CADD-Splice-improving genome-wide variant effect prediction using deep learning-derived splice scores. Genome Med. 13, 31 (2021).

58. X. Liu, C. Li, C. Mou, Y. Dong, Y. Tu, dbNSFP v4: a comprehensive database of transcript-specific functional predictions and annotations for human nonsynonymous and splice-site SNVs. Genome Med. 12, 103 (2020).

59. L. A. Miosge, et al., Comparison of predicted and actual consequences of missense mutations. Proc Natl Acad Sci USA 112, E5189–5198 (2015).

60. R. Ghosh, N. Oak, S. E. Plon, Evaluation of in silico algorithms for use with ACMG/AMP clinical variant interpretation guidelines. Genome Biol. 18, 225 (2017).

61. N. Whiffin, et al., Using high-resolution variant frequencies to empower clinical genome interpretation. Genet. Med. 19, 1151–1158 (2017).

62. Z. Liu, L. Zhu, R. Roberts, W. Tong, Toward Clinical Implementation of Next-Generation Sequencing-Based Genetic Testing in Rare Diseases: Where Are We? Trends Genet. 35, 852–867 (2019).

63. H. Nakagawa, M. Fujita, Whole genome sequencing analysis for cancer genomics and precision medicine. Cancer Sci. 109, 513–522 (2018).

64. A. H. D. M. Dam, et al., Homozygous mutation in SPATA16 is associated with male infertility in human globozoospermia. Am. J. Hum. Genet. 81, 813–820 (2007).

65. M. Gershoni, et al., A familial study of azoospermic men identifies three novel causative mutations in three new human azoospermia genes. Genet. Med. 19, 998–1006 (2017).

66. Y. Fujihara, A. Oji, T. Larasati, K. Kojima-Kita, M. Ikawa, Human Globozoospermia-Related Gene Spata16 Is Required for Sperm Formation Revealed by CRISPR/Cas9-Mediated Mouse Models. Int. J. Mol. Sci. 18 (2017).

67. R. Guo, et al., The ssDNA-binding protein MEIOB acts as a dosage-sensitive regulator of meiotic recombination. Nucleic Acids Res. 48, 12219–12233 (2020).

68. B. Mandon-Pépin, et al., Genetic investigation of four meiotic genes in women with premature ovarian failure. Eur. J. Endocrinol. 158, 107–115 (2008).

69. J. Hikiba, et al., Structural and functional analyses of the DMC1-M200V polymorphism found in the human population. Nucleic Acids Res. 36, 4181–4190 (2008).

70. Q. Li, et al., CRISPR-Cas9-mediated base-editing screening in mice identifies DND1 amino acids that are critical for primordial germ cell development. Nat. Cell Biol. 20, 1315–1325 (2018).

71. S. Özkan, et al., “The computational approach to variant interpretation” in Clinical DNA Variant Interpretation, (Elsevier, 2021), pp. 89–119.

72. S. N. Kobren, et al., Commonalities across computational workflows for uncovering explanatory variants in undiagnosed cases. Genet. Med. 23, 1075–1085 (2021).

73. P. D. Stenson, et al., The Human Gene Mutation Database: towards a comprehensive repository of inherited mutation data for medical research, genetic diagnosis and next-generation sequencing studies. Hum. Genet. 136, 665–677 (2017).

74. D. G. Grimm, et al., The evaluation of tools used to predict the impact of missense variants is hindered by two types of circularity. Hum. Mutat. 36, 513–523 (2015).

75. N. Shah, et al., Identification of misclassified clinvar variants via disease population prevalence. Am. J. Hum. Genet. 102, 609–619 (2018).

76. P. Kumar, S. Henikoff, P. C. Ng, Predicting the effects of coding non-synonymous variants on protein function using the SIFT algorithm. Nat. Protoc. 4, 1073–1081 (2009).

77. I. A. Adzhubei, et al., A method and server for predicting damaging missense mutations. Nat. Methods 7, 248–249 (2010).

78. J. Ren, et al., DOG 1.0: illustrator of protein domain structures. Cell Res. 19, 271–273 (2009).

79. J. J. Hardy, et al., Variants in GCNA, X-linked germ-cell genome integrity gene, identified in men with primary spermatogenic failure. Hum. Genet. 140, 1169–1182 (2021).

80. P. N. Schlegel, et al., Diagnosis and treatment of infertility in men: AUA/ASRM guideline part I. Fertil. Steril. 115, 54–61 (2021).

81. L. Björndahl, J. Kirkman Brown, other Editorial Board Members of the WHO Laboratory Manual for the Examination and Processing of Human Semen, The sixth edition of the WHO Laboratory Manual for the Examination and Processing of Human Semen: ensuring quality and standardization in basic examination of human ejaculates. Fertil. Steril. 117, 246–251 (2022).

82. L. Nagirnaja, et al., Diverse monogenic subforms of human spermatogenic failure. medRxiv (2022) https://doi.org/10.1101/2022.07.19.22271581.

83. G. E. Truett, et al., Preparation of PCR-quality mouse genomic DNA with hot sodium hydroxide and tris (HotSHOT). BioTechniques 29, 52–54 (2000).

84. E. Bolcun-Filas, V. D. Rinaldi, M. E. White, J. C. Schimenti, Reversal of female infertility by Chk2 ablation reveals the oocyte DNA damage checkpoint pathway. Science 343, 533–536 (2014).

85. A. H. Peters, A. W. Plug, M. J. van Vugt, P. de Boer, A drying-down technique for the spreading of mammalian meiocytes from the male and female germline. Chromosome Res. 5, 66–68 (1997).

86. L. Uroz, O. Rajmil, C. Templado, Premature separation of sister chromatids in human male meiosis. Hum. Reprod. 23, 982–987 (2008).

87. A. Henrie, et al., ClinVar Miner: Demonstrating utility of a Web-based tool for viewing and filtering ClinVar data. Hum. Mutat. 39, 1051–1060 (2018).

