## Supplemental Figs 1-11 for "*In vivo* versus *in silico* assessment of potentially pathogenic missense variants in human reproductive genes"

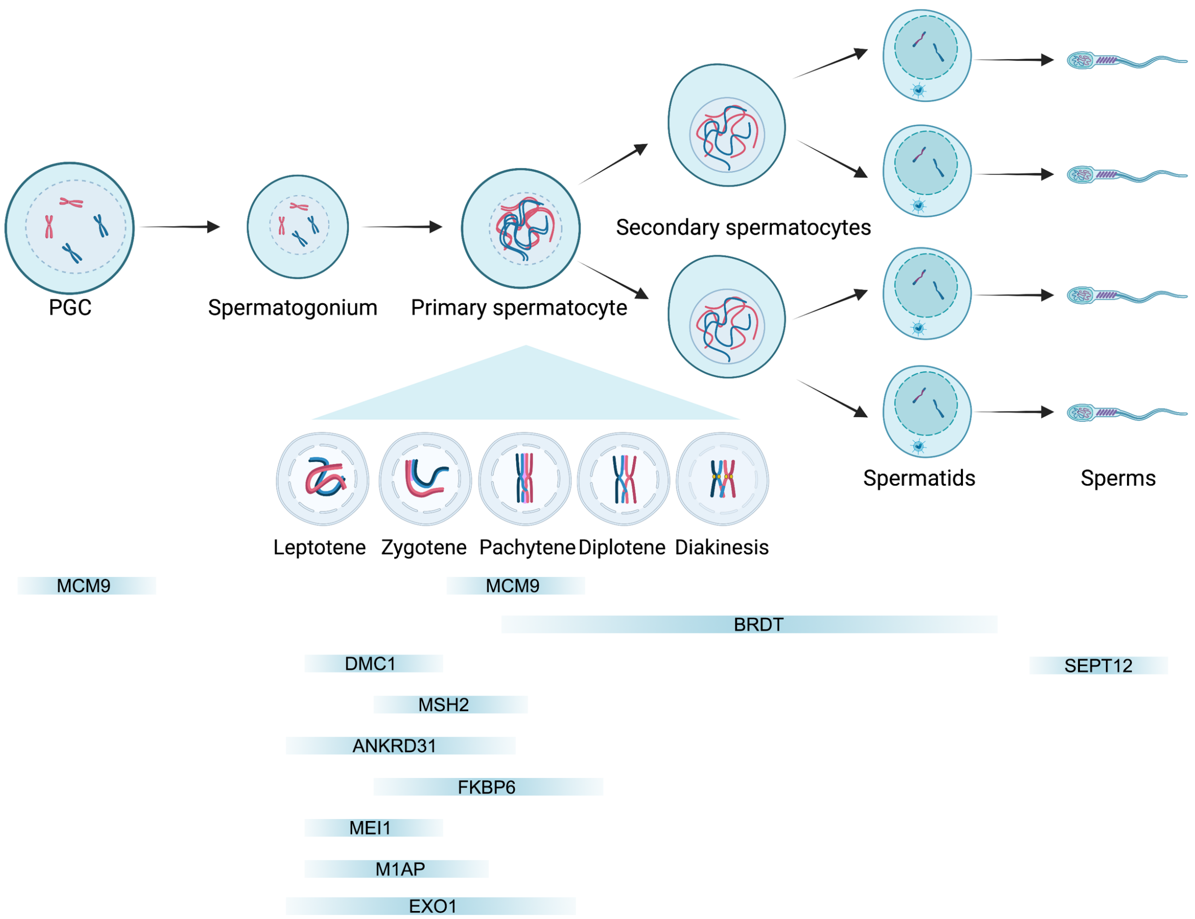


**Figure S1. Spermatogenesis schematic showing temporal involvement of ten studied genes.** The light blue horizontal bars represent the gametogenesis stage that is impacted in knockout mice.

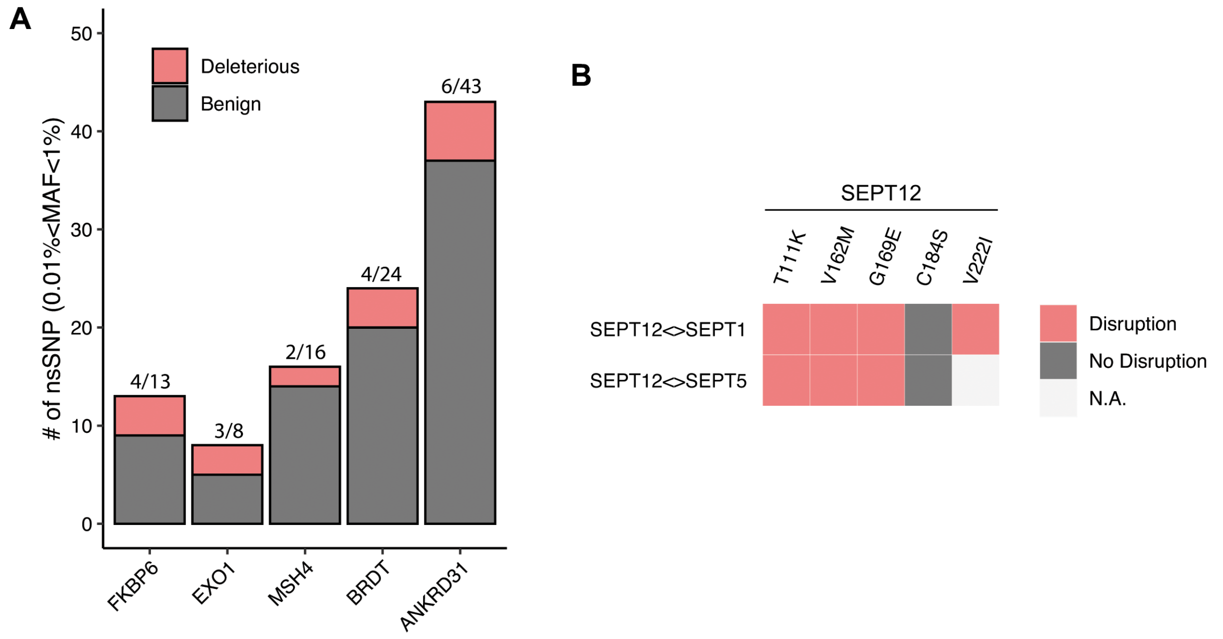


**Figure S2. Selection of causal variants from reproduction genes.** A) Distribution of benign and deleterious missense variants (0.01%<MAF<1%) predicted by SIFT and PolyPhen2 in indicated reproduction genes. B) Y2H screening of missense variants in SEPT12 disrupting its interaction with the SEPT1 and SEPT5 proteins, respectively. N.A., not analyzed.

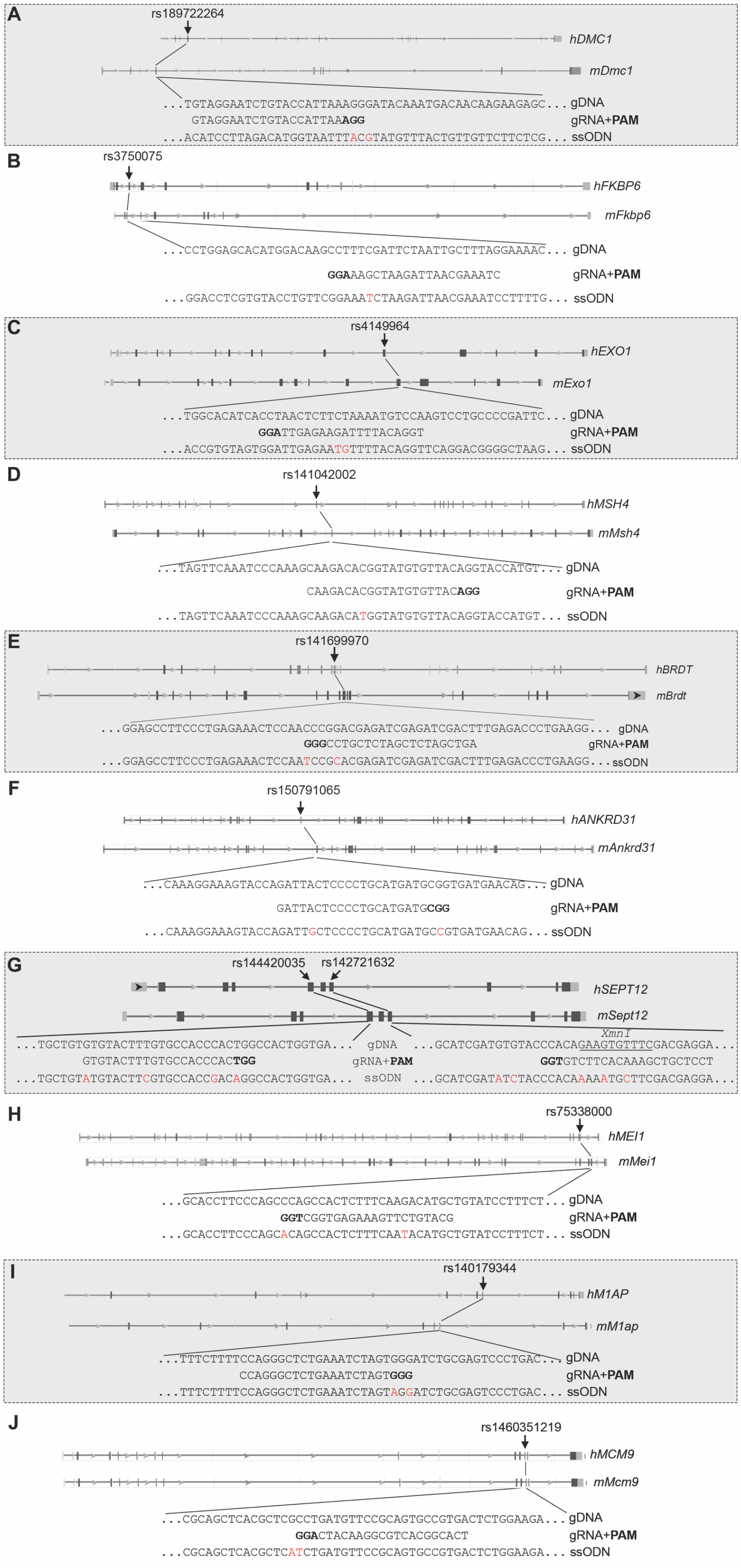


**Figure S3. CRISPR-Cas9 editing strategies.** PAM sites are in bold. The red fonts in ssODNs represent the edited version of targeted nucleotides.

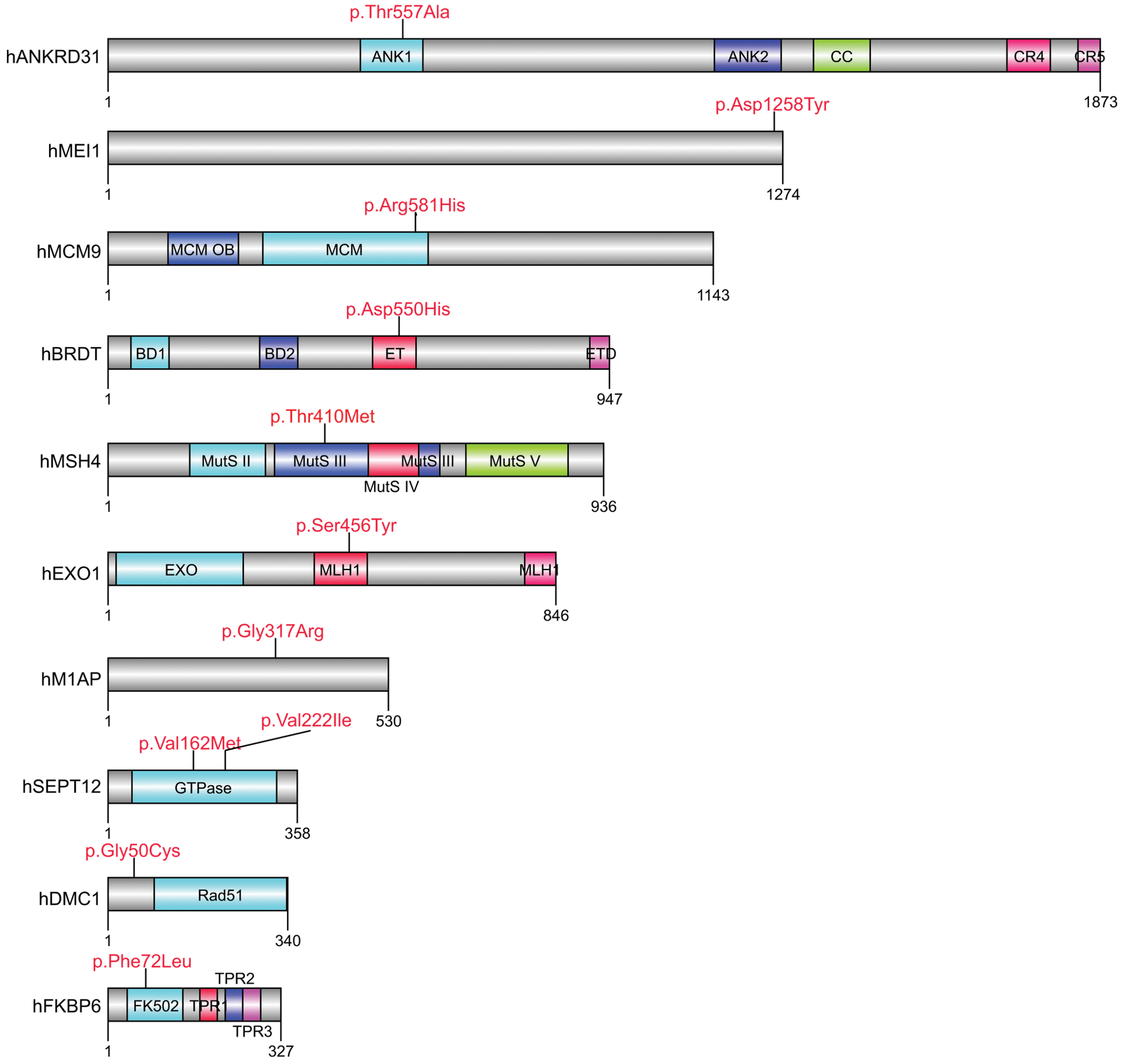


**Figure S4. Schematic of human protein domain structures and variant locations.**

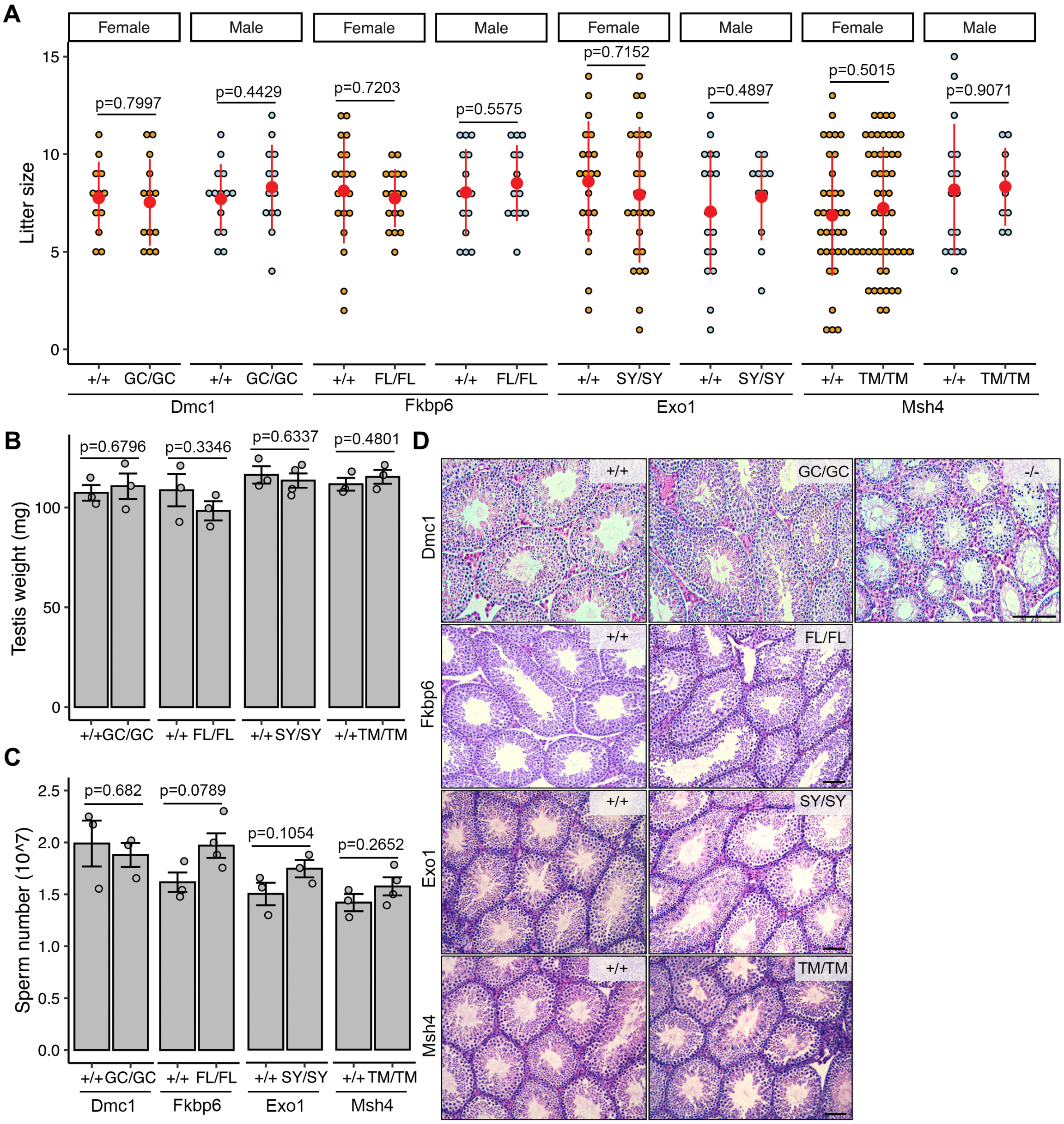


**Figure S5. Phenotypic characterization of homozygous mice harboring missense variants in *DMC1*, *MSH4*, *EXO1* and *FKBP6*.** A) Litter sizes from mating of *Dmc1*$^{G50C/G50C}$ (GC/GC), *Fkbp6*$^{F72L/F72L}$ (FL/FL), *Exo1*$^{S465Y/S465Y}$ (SY/SY) and *Msh4*$^{T410M/T410M}$ (TM/TM) females and males to WT partners. N=2 for GC/GC, FL/FL and SY/SY male and female mice, N=3 for TM/TM male and female mice. B) Testis weights of 2-month old mice. C) Sperm counts in mutant homozygotes and littermate controls.  D) Cross-sections of testes showing comparable histology to the littermate controls. Scale bars = 100 μm. Data in A, B and C are represented as the mean ± SEM and were analyzed using two-tailed unpaired *t test*.

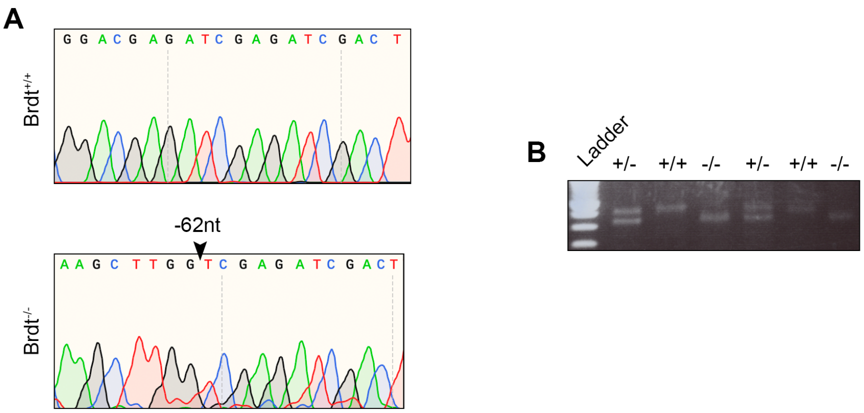


**Figure S6. Genotyping *Brdt*$^{-/-}$ mice.** A) Representative Sanger sequencing chromatograms from WT and mutant mice. Black arrowhead above the chromatogram indicates the breakpoint of the 62nt deletion. B) Representative PCR genotyping results of pups from a het X het mating.

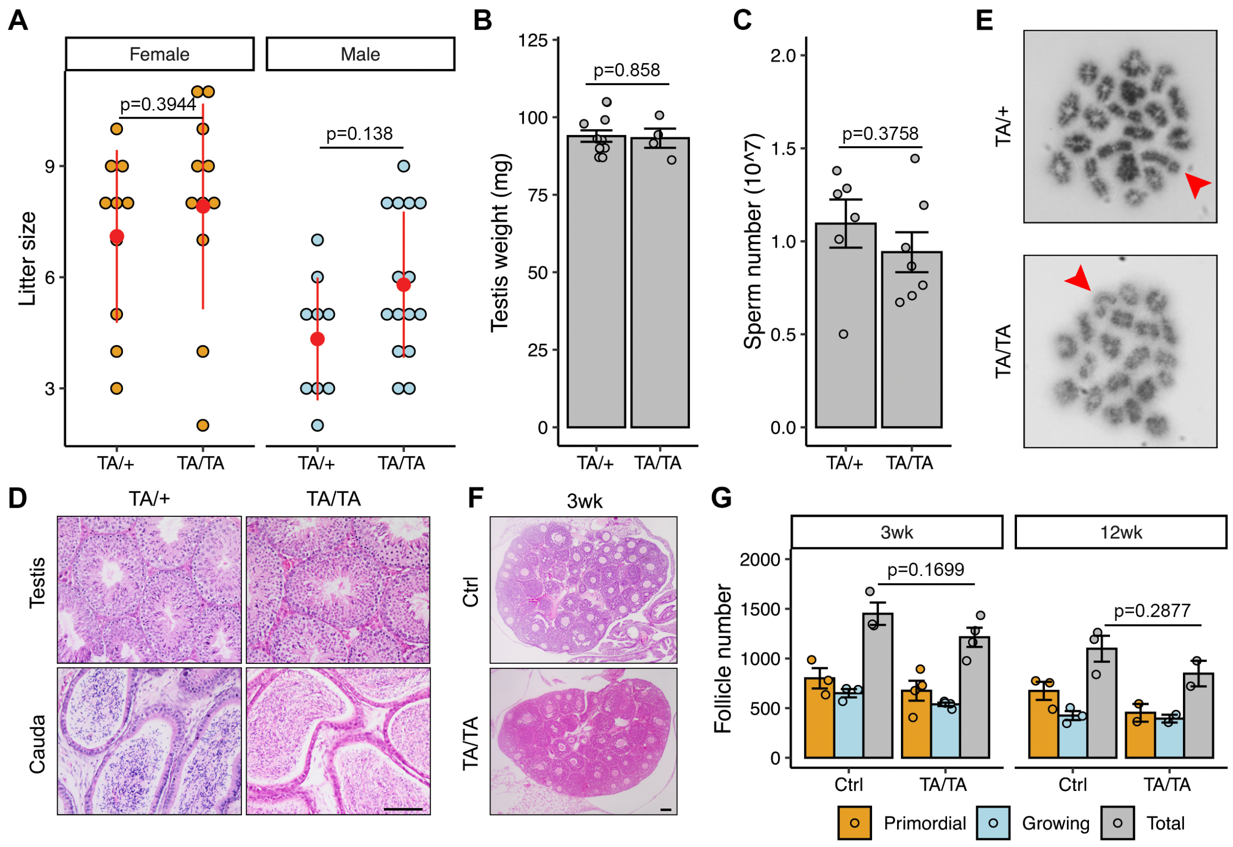


**Figure S7. Phenotypic analysis of *Ankrd31* p.T557A mice.** A) Litter sizes from mating of *Ankrd31*$^{T557A/T557A}$ (TA/TA) and *Ankrd31*$^{T557A/+}$ (TA/+) animals to WT partners. N=3 for TA/TA male and female mice. B) Testis weights of 2-month-old mice. C) Sperm counts. D) Histological analyses of 2-month old testes and cauda epididymides. Scale bars = 100 µm. E) Meiotic metaphase I spermatocyte chromosome preparations. Arrows indicate X and Y chromosomes. F) Histological analyses of 2-month old ovaries. Scale bars = 100 µm. G) Follicle counts summed across every fifth serial section. Data in A, B, C and G are represented as the mean ± SEM and were analyzed using two-tailed unpaired *t test*.

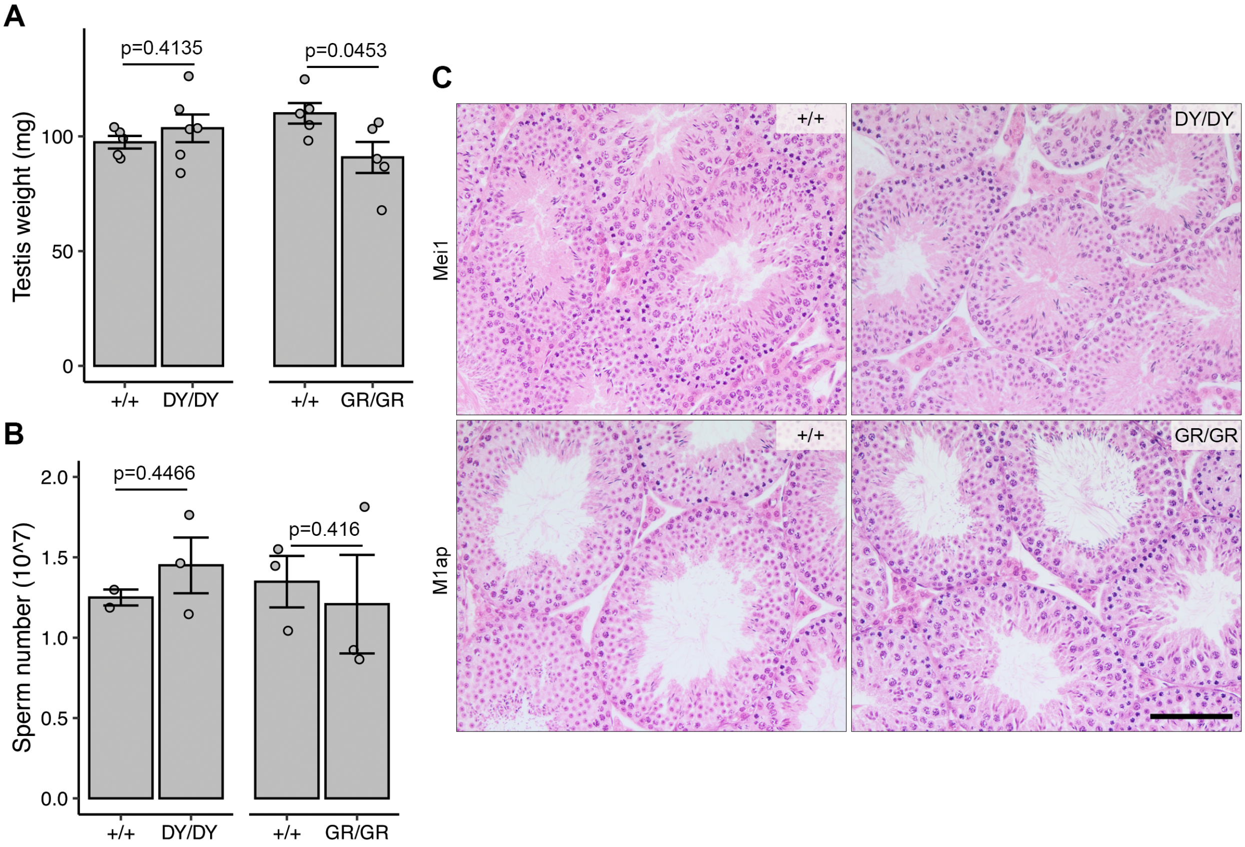


**Figure S8. Phenotypic analysis of *Mei1* p.D1258Y and *M1ap* p.G317R mice.** A) Testis weights of 2-month old mice. B) Sperm counts. C) Histological analyses of 2-month old testes. Scale bars = 100 µm. Data in A and B are represented as the mean ± SEM and were analyzed using two-tailed unpaired *t test*.

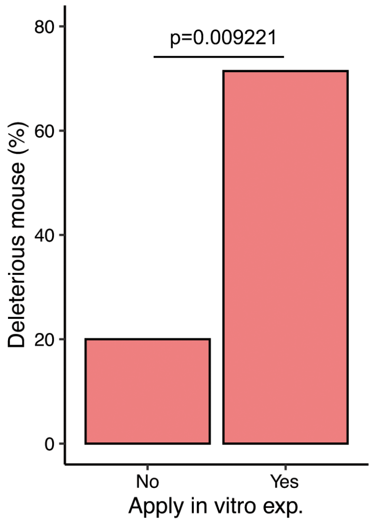


**Figure S9. *In vitro* pre-screen experiments can improve the selection of variants that yield phenotypes in mouse models.** The *p* value was calculated by two-sided Fisher's Exact Test.

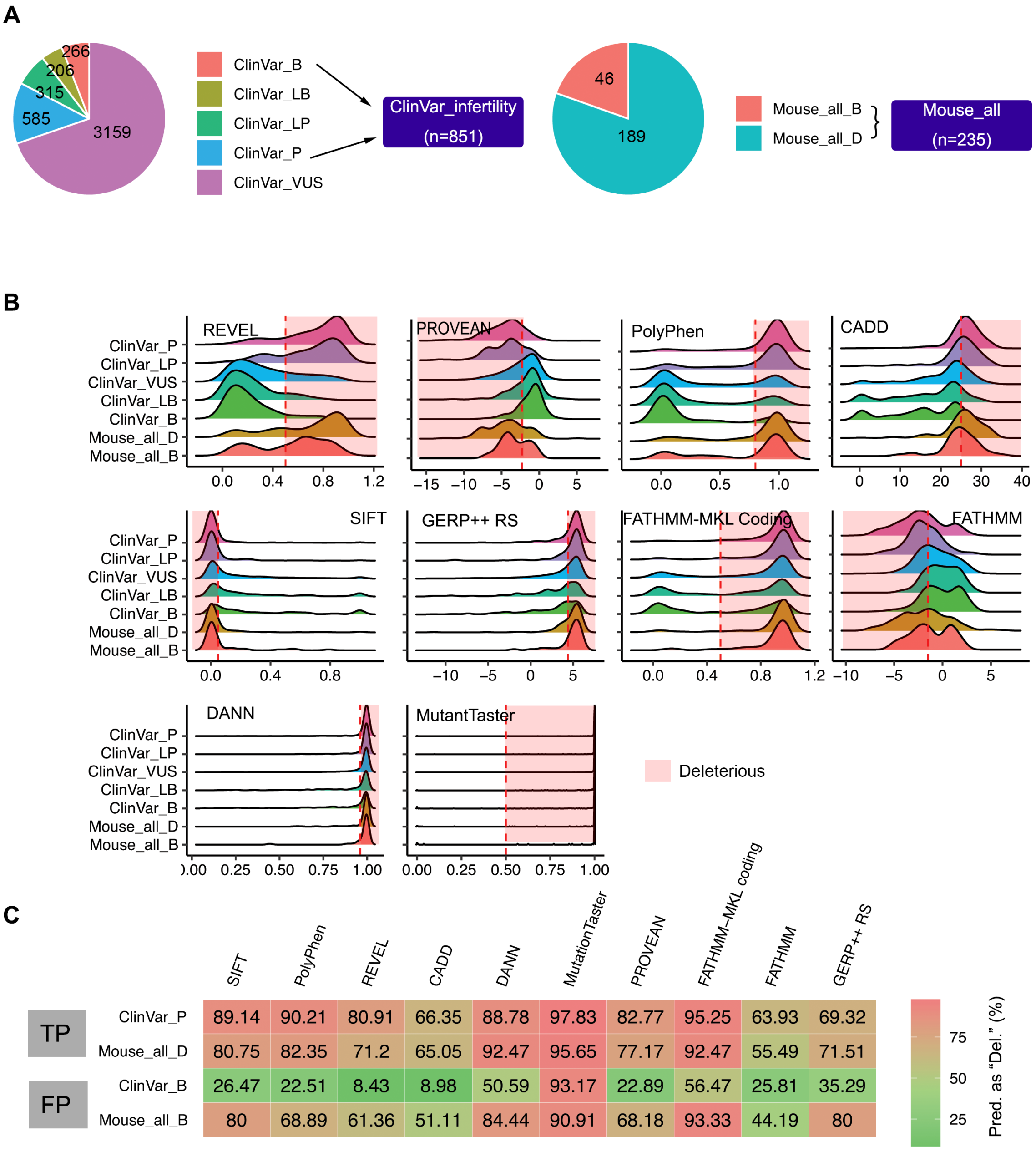


**Figure S10. Prediction of ClinVar and mouse-modeled missense variants.** A) Distribution of missense variants in two datasets. The variant numbers are indicated. B) Distribution of prediction scores by 10 algorithms. The red dotted line represents the cutoff, and the pink rectangle highlights the deleteriousness scores. C) Percentage of variants predicted as deleterious by indicated algorithms from the indicated catalogs (in rows). TP = true positives; FP = false positives; Del and D = deleterious; B = benign; LB = likely benign; VUS = variants of uncertain significance; P = pathogenic; LP = likely pathogenic.

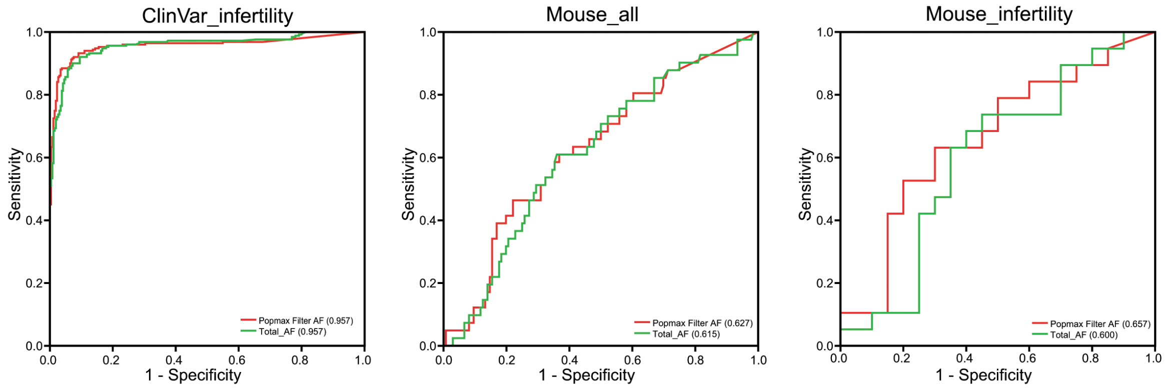


**Figure S11. ROC curves of Popmax filtering AF and Total AF in three datasets.** Logarithmic values were plotted. AUC values close to 1 indicate deleterious/pathogenic variants were clearly separated from benign variants, whereas scores close to 0.5 suggests that the variants were distributed randomly.
